# NetMix2: Unifying network propagation and altered subnetworks

**DOI:** 10.1101/2022.01.31.478575

**Authors:** Uthsav Chitra, Tae Yoon Park, Benjamin J. Raphael

**Affiliations:** Department of Computer Science, Princeton University; Lewis-Sigler Institute for Integrative Genomics, Princeton University

## Abstract

A standard paradigm in computational biology is to use interaction networks to analyze high-throughput biological data. Two common approaches for leveraging interaction networks are: (1) *network ranking*, where one ranks vertices in the network according to both vertex scores and network topology; (2) *altered subnetwork* identification, where one identifies one or more subnetworks in an interaction network using both vertex scores and network topology. The dominant approach in network ranking is network propagation which smooths vertex scores over the network using a random walk or diffusion process, thus utilizing the global structure of the network. For altered subnetwork identification, existing algorithms either restrict solutions to subnetworks in *subnetwork families* with simple topological constraints, such as connected subnetworks, or utilize ad hoc heuristics that lack a rigorous statistical foundation. In this work, we unify the network propagation and altered subnetwork approaches. We derive a subnetwork family which we call the *propagation family* that approximates the subnetworks ranked highly by network propagation. We introduce NetMix2, a principled algorithm for identifying altered subnetworks from a wide range of subnetwork families, including the propagation family, thus combining the advantages of the network propagation and altered subnetwork approaches. We show that NetMix2 outperforms network propagation on data simulated using the propagation family. Furthermore, NetMix2 outperforms other methods at recovering known disease genes in pan-cancer somatic mutation data and in genome-wide association data from multiple human diseases. NetMix2 is publicly available at https://github.com/raphael-group/netmix2.

## 1 Introduction

Biological systems consist of interactions between many components. These interactions are often represented with networks, e.g., protein-protein interaction networks or gene regulatory networks. A standard paradigm in computational biology is to use an interaction network as prior knowledge for interpreting high-throughput, genome-scale data. Interaction networks have informed the analysis of biological data in many different applications including protein function prediction [1, 2, 3, 4, 5], differential expression analysis [6, 7, 8, 9, 10, 11, 12], prioritization of germline variants [13, 14, 15, 16, 17, 18], identification of driver mutations in cancer [19, 20, 21, 22, 23, 24], and more [25, 26, 13, 27, 28, 29, 30, 31, 31, 32, 33, 34, 35].

Numerous methods that use interaction networks in interpreting high-throughput ‘omics data have been developed (reviewed in [28, 36, 37, 22, 38, 39]). While the algorithmic details of these methods are diverse, nearly all of them employ one of two different strategies. The first strategy is *network ranking*, where one is given either a subset of vertices (genes/proteins) or a score for each vertex (gene/protein), and the goal is to rank all vertices according to both the subset/scores and the positions of vertices in the network. Early network ranking algorithms relied on the “guilt-by-association” principle, or the idea that genes/proteins with similar functions are directly connected in the interaction network. These “direct connection” algorithms were typically applied in *semi-supervised* settings, e.g., protein function prediction or disease-gene prioritization [40, 41], where only a subset of vertices are known to have a specific biological function. Later, inspired by the success of random walk, diffusion, and graph kernel methods in statistics and machine learning (e.g., the PageRank algorithm [42]), *network propagation* — also known as label propagation [43] — became the dominant approach for network ranking [38]. Briefly, network propagation involves using a random walk or diffusion process to “smooth” vertex scores across a network. Following [38], we use the term network propagation to refer to the broad class of methods that smooth scores over a network using a random walk or diffusion process. This includes popular processes like the random walk with restart (i.e., PageRank) [42], but also other processes including diffusion state distance [44, 45] or the heat kernel [20, 46]. By using these random walk/diffusion processes, network propagation methods simultaneously account for all possible paths between vertices. Thus, in contrast to the early methods which only use “direct connections” (edges) between vertices, network propagation methods fully utilize the *global* structure of the network. Indeed, network propagation has even been shown to be asymptotically optimal for network ranking for some random graph models [47].

The second strategy is the identification of *altered subnetworks*, also called *network modules* or *active subnetworks*. Here, the input is a measurement or a score for each vertex of the interaction network (e.g., *p*-values from differential gene expression), and the goal is to identify subnetworks (modules) that contain high scoring vertices that are “close” in the network^1^. Altered subnetwork approaches rely on the specification of a *subnetwork family*, or a family of possible subnetworks; sometimes the family is stated explicitly – e.g., the early approaches such as jActiveModules [49] or heinz [8] identify connected subnetworks — but in other methods, the subnetwork family is implicitly specified — e.g., the optimization problems of [50] and [51] penalize subnetworks with large cut-size and small edge-density, respectively. The altered subnetwork approach is closely related to the identification of *network anomalies* in the data mining and machine learning literature [52, 53, 54, 55, 56, 57, 58]. In contrast to ranking algorithms that yield a ranking of all vertices, altered subnetwork approaches output one or more *subsets* of vertices, and thus explicitly estimate the number of vertices in the altered subnetworks. A major challenge with altered subnetwork approaches is to choose an appropriate subnetwork family. For example, connectivity is often too weak of an assumption for biological networks, e.g., some methods that use connectivity identify large subnetworks [59] because of a statistical bias in a commonly used test statistic [60, 61].

There have been a few attempts to bridge the gap between network propagation approaches and altered subnetwork approaches, combining the modeling of global network topology from network propagation with the optimization over subnetwork families from altered subnetwork approaches. For example, PRINCE [62] first propagates the vertex scores and identifies altered subnetworks as edge-dense subnetworks whose vertices have large network propagated scores. The HotNet algorithms [20, 63, 23, 64] identify altered subnetworks by finding clusters in a weighted and directed graph derived from network propagation. TieDIE [65] propagates two sets of vertex scores and aims to find high-scoring subnetworks for both sets of propagated scores. More recently, the NetCore algorithm [66] finds subnetworks whose vertices have large node “coreness” and large propagated scores. However, none of these approaches give an explicit definition of the subnetwork family, instead relying on heuristics to identify the altered subnetwork after performing network propagation. Because of these heuristics, network propagation approaches typically do not have provable guarantees for altered subnetwork identification. In contrast, methods that explicitly rely on a well-defined subnetwork family often have statistical or theoretical guarantees, e.g., jActiveModules [49] computes a maximum likelihood estimator while our recent estimator NetMix is asymptotically unbiased [60, 61].

Another practical issue is the evaluation of altered subnetwork methods. Most network algorithms demonstrate their performance by benchmarking their algorithm against existing network algorithms. While these comparisons are useful, they may also hide biases shared between algorithms. For example, Lazareva et al. [67] observed that some well-known network algorithms have a bias towards high-degree vertices in the interaction network, while Levi et al. [68] similarly observed a bias in GO term enrichment among well-known network algorithms. In order to quantify the potential biases of altered subnetwork algorithms, these algorithms need to be compared against carefully selected baselines, including baselines that do not use the interaction network and baselines that do not use the vertex scores.

In this paper, we introduce NetMix2, an algorithm which unifies the network propagation and altered subnetwork approaches. NetMix2 generalizes NetMix [60] to a wide range of subnetwork families and vertex score distributions. Specifically, NetMix2 takes as input a wide variety of subnetwork families, including not only the “connected family” used in existing altered subnetwork methods [6, 8, 60] but also any subnetwork family defined by linear or quadratic constraints, such as subnetworks with high edge density or subnetworks with small cut-size. We use this flexibility to investigate the topology of subnetworks identified by network propagation methods. We show empirically that network propagation does not correspond to standard topological constraints on altered subnetworks such as connectivity [8, 6, 60], cut-size [50], or edge-density [51]. Instead, we derive the *propagation family*, a subnetwork family that we show “approximates” the subnetworks identified by network propagation approaches and thereby unifies the two major network approaches in the literature: network propagation and altered subnetwork identification. NetMix2 also uses local false discovery rate (local FDR) methods [69, 70, 71] to flexibly model vertex score distributions, in contrast to the strict parametric assumptions made by existing methods [8, 60].

On simulated data we show that NetMix2 outperforms network propagation for subnetworks from the propagation family and other common subnetwork families. Interestingly, NetMix2 outperforms network propagation by the largest margin for the propagation family. We then apply NetMix2 with the propagation family to cancer mutation data and genome-wide association studies (GWAS) data from several complex diseases. On cancer data, we show that NetMix2 outperforms existing network propagation and altered subnetwork methods in identifying cancer driver genes. On GWAS data, we demonstrate that network propagation often has similar performance to simple baselines that only use the vertex scores or only use the network. However, in cases where network propagation outperforms these baselines, we show that NetMix2 outperforms network propagation.

## 2 Methods

### 2.1 Altered Subnetwork Problem

We start by formalizing the problem of altered subnetwork identification. Let *G* = (*V, E*) be an interaction network with a score *X*_*v*_ for each vertex *v*. We assume there is an *altered subnetwork A ⊆ V* whose scores {*X*_*v*_} _*v*∈*A*_ are drawn independently from a different distribution than the scores {*X*_*v*_}_*v* ∉*A*_ of vertices not in the altered subnetwork *A*. The topology of the altered subnetwork is described by membership in a *subnetwork family 𝒮* ⊆ 𝒫(*V*), where 𝒫(*V*) denotes the power set of all subsets of vertices *V*.

Following the exposition in [60, 61], we model the distribution of the scores **X** = {*X*_*v*_}_*v*∈*V*_ as the Altered Subnetwork Distribution (ASD).

#### Altered Subnetwork Distribution (ASD)

*Let G* = (*V, E*) *be a graph, let 𝒮* ⊆ 𝒫 (*V*) *be a subnetwork family, and let A* ∈ 𝒮. *We say* **X** = (*X*_*v*_)_*v*∈*V*_ *is distributed according to the* Altered Subnetwork Distribution *ASD*_S_(*A, 𝒟*_*a*_, *𝒟*_*b*_) *provided the X*_*v*_ *are independently distributed as*

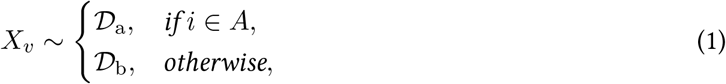

*where 𝒟*_a_ *is the* altered *distribution and 𝒟*_b_ *is the* background *distribution*.

The distribution ASD_S_(*A, 𝒟* _*a*_, *𝒟*_*b*_) is parameterized by four quantities: the altered subnetwork *A*, the subnetwork family 𝒮, the altered distribution 𝒟_a_, and the background distribution 𝒟_b_.

Given the measurements **X** ∼ ASD_𝒮_ (*A, 𝒟*_*a*_, *𝒟*_*b*_) and the subnetwork family 𝒮 ⊆ 𝒫 (*V*), the goal of the *Altered Subnetwork Problem* is to identify the altered subnetwork *A*. We formalize this problem below.

#### Altered Subnetwork Problem (ASP)

*Given* **X** ∼ ASD _𝒮_ (*A, 𝒟*_*a*_, *𝒟*_*b*_) *and subnetwork family 𝒮, find A*.

The ASP describes a broad class of problems that are studied in many fields including computational biology [8, 6, 60], statistics [72, 54, 57, 73] and machine learning [55, 61, 74], with different problems making different choices for (1) the distributions 𝒟_a_, 𝒟_b_ and (2) the subnetwork family 𝒮. Two prominent examples of distributions 𝒟_a_, 𝒟_b_ that have been previously studied in the biological literature are the following.

- **Normal distributions:** 𝒟_a_ = *N*(*μ*, 1) and 𝒟_b_ = *N* (0, 1). Normal distributions are often used to model z-scores [75, 76, 60, 77, 78]. We call the ASP and ASD with these distributions the *normally distributed ASP* and *normally distributed ASD*, respectively; for notational convenience, we use NASD _𝒮_ (*A, μ*) to refer to the normally distributed ASD.
- **Beta-uniform distributions:** 𝒟_a_ = Beta(*a*, 1) and 𝒟_b_ = Uni(0, 1). Beta-uniform mixture distributions are another common model for p-value distributions [8, 79]. We call the ASP, ASD with these distributions the *Beta-Uniform ASP* and *Beta-Uniform ASD*, respectively.

We also list several examples of subnetwork families 𝒮, with each subnetwork family corresponding to a different topological assumption on the altered subnetwork *A*. Some of these families have been explicitly applied in biological settings, while other families formalize topological constraints that are implicitly made in the biological literature.

- 𝒮 = 𝒞_*G*_, the *connected family*, or the set of all connected subgraphs *S* of an interaction network *G*. [60, 6, 8] identify altered subnetworks by solving the ASP for the connected family 𝒞_*G*_.
- 𝒮 = *ℰ*_*G,p*_, the *edge-dense family*, or the set of all subgraphs *S* of *G* with edge-density 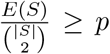, where *E*(*S*) = |{(*u, v*) ∈*E* : *u* ∈ *S, v* ∈ *S*}| is the number of edges between vertices in *S*. The edge-dense family *ℰ*_*G,p*_ formalizes the topological constraints made by [51, 80, 62], which identify altered subnetworks that have large edge-density.
- 𝒮 = 𝒯_*G,ρ*_, the *cut family*, or the set of all subgraphs *S* of *G* with 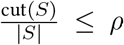, where cut(*S*) =|{(*u, v*) ∈*E* : *u* ∈*S, v* ∉*S* is the number of edges with exactly one endpoint in *S*. The cut family 𝒯_*G,ρ*_ formalizes the topological constraints made by [81], which identifies altered subnetworks that have small cut.
- 𝒮 = 𝒬_*G,ρ*_, the *modularity family*, or the set of all subgraphs *S* of *G* with modularity *Q*(*S*) ≥ *ρ*. The modularity family formalizes the topological constraints made by [82], which identifies altered subnetworks that have high modularity.

We note that the ASP — with the subnetwork families 𝒮 described above — describes the problem of identifying a single altered subnetwork in a network *G*. By creating a new subnetwork family consisting of the union of *k* disjoint subnetworks in family 𝒮, the ASP also describes the problem of identifying *multiple* altered subnetworks.

Early methods for identifying altered subnetwork solved the ASP for the connected family 𝒮 = 𝒞_*G*_ and different choices of vertex score distributions 𝒟_a_, 𝒟_b_. For example, two seminal methods, jActiveModules [6] and heinz [8], solve the normally distributed and Beta-Uniform ASP, respectively, with the connected family 𝒮= 𝒞_*G*_. Recently we showed that many existing methods, including jActiveModules and heinz, are *biased*, in the sense that they typically estimate subnetworks *A* that *Â* are much larger than the altered subnetwork *A* [60, 61]. To this end, we derived the NetMix algorithm, which finds an asymptotically unbiased *Â*_NetMix_ of the altered subnetwork *A* for the connected family 𝒮 = 𝒞 _*G*_. However, as we demonstrate in [60] and Section 3 below, many of these methods — including NetMix — have comparable performance to a naive “scores-only” baseline that does not use the network *G*.

### 2.2 Network Propagation and the Propagation Family

Another strategy often used to incorporate interaction networks *G* with high throughput biological data is *network propagation*. Network propagation involves the use of random walk or diffusion processes to “smooth” or “propagate” vertex scores *X*_*v*_ across a network [38]. Formally, given vertex scores *X*_*v*_, the *network propagated scores Y*_*v*_ are computed as

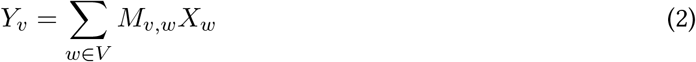

where *M* ∈ ℝ^|*V* |×|*V* |^ is a similarity matrix on the vertices *V* of the network *G* typically derived from a random walk on *G*. One popular choice for the similarity matrix *M* is the random walk with restart (personalized PageRank) similarity matrix *M*_PPR_ = *r*(*I*− (1− *r*)*P*)^−1^, where *r* ∈ (0, 1) is the restart probability, *I* is the identity matrix, and *P* is the transition matrix for a random walk with restart on *G*.

A few methods have attempted to use network propagation to identify the altered subnetwork *A* from propagated scores *Y*_*v*_, e.g., PRINCE [62] finds edge-dense subnetwork with large propagated scores *Y*_*v*_. These methods implicitly assume that the propagated scores *Y*_*v*_ are larger for vertices *v* ∈*A* in the altered subnetwork *A* compared to vertices *v* ∉*A* not in the altered subnetwork *A* ∈ 𝒮 However, we empirically find (Section 3.1) that this assumption generally does not hold for altered subnetworks *A* from the connected family 𝒮= 𝒞_*G*_, the edge-dense family 𝒮= *ℰ*_*G,p*_ and the cut family 𝒮 =𝒯_*G,ρ*_, which suggests that network propagation methods do not solve the ASP with these subnetwork families 𝒮.

Thus, we derive a subnetwork family 𝒮 that approximates the subnetworks identified by network propagation methods. Informally, we first note that network propagation methods identify altered subnetworks *A* whose vertices *v* ∈ *A* have large propagated scores *Y*_*v*_. We observe that — by making the simplifying assumption that the vertex scores *X*_*v*_ = 1 {_*v* ∈*A*}_ are binary — the propagated score *Y*_*v*_ = Σ_*w* ∈*A*_ *M*_*v,w*_ of a vertex *v* is large if the similarities *M*_*v,w*_ is large for many *w* ∈ *A*. Intuitively, one natural way to enforce that the similarities *M*_*v,w*_ are large is to lower-bound them, i.e., require that *M*_*v,w*_ ≥ *δ* for many *w* ∈*A* and for some (large) constant *δ >* 0.

This intuition motivates the formal definition of the *propagation family ℳ*_*δ,p*_, or the set of all subgraphs *S* with *M*_*u,v*_≥ *δ* and *M*_*v,u*_ ≥ *δ* for *p* fraction of tuples (*u, v*) ∈ *S*. (Because the similarity matrix *M* may not be symmetric, we constrain both *M*_*u,v*_ and *M*_*v,u*_.) We note that the propagation family ℳ_*δ,p*_ is equal to the edge-dense family 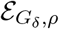 for the similarity threshold graph *G*_*δ*_ = (*V, E*_*δ*_), which has edge (*u, v*) ∈ *E*_*δ*_ if and only if *M*_*u,v*_ ≥ *δ* and *M*_*v,u*_ ≥ *δ*.

We partially formalize our derivation of the propagation family ℳ_*δ,p*_ through the following result, which bounds the false discovery rate (FDR) of the subnetwork consisting of the largest propagated scores for data 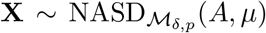 from the normally distributed ASD with propagation family ℳ_*δ,p*_ and with density *p* = 1, i.e., cliques in the similarity threshold graph *G*_*δ*_.

#### Proposition 1.

*Let G = (V,E) be a graph and let M* ∈ [0,1] ^|*V* |×|*V* |^ *be a matrix indexed by vertices V*. *Define r* = min_*v*∈*V*_ *M*_*v,v*_, *c* = max_*v*∉*V*_ Σ_*w*∈*A*_ *M*_*v,w*_, *and d* = max_*v,w*∈*V*_ Σ_*u*∈*V*_ (*M*_*w,u*_ − *M*_*v,u*_)^2^. *Let* 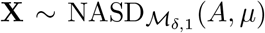 *where μ* ≥ 1 *and* 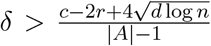. *Then with probability at least* 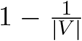, *the altered subnetwork A consists of the* |*A*| *vertices with the largest propagated scores Y*_*v*_.

### 2.3 NetMix2

We derive the NetMix2 algorithm, which solves the ASP for a wide range of subnetwork families 𝒮 and distributions 𝒟_a_, 𝒟_b_ (Figure 1). In particular, NetMix2 solves the ASP for the propagation family ℳ_*δ,p*_, and thus bridges the gap between the ASP and network propagation.

**Figure 1:**
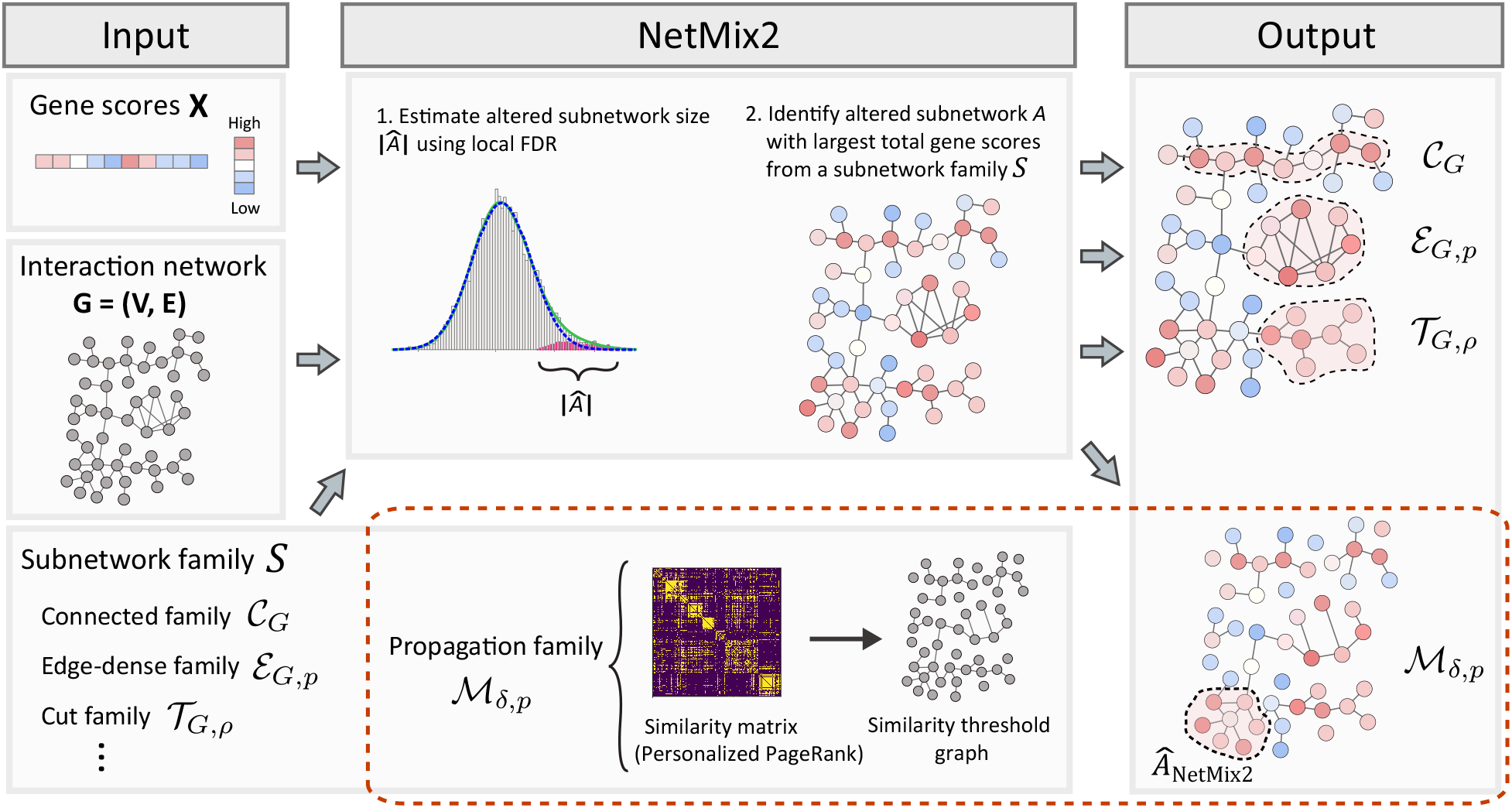
Overview of the NetMix2 algorithm. The inputs to NetMix2 are a graph *G*, gene scores {*X*_*v*_}_*v* ∈*V*_, and a subnetwork family 𝒮. First, NetMix2 computes an estimate |*Â*| of the size |*A*| of the altered subnetwork *A* using local false discovery rate (local FDR). Next, NetMix2 solves an optimization problem to identify the subnetwork *S* ∈ 𝒮 size |*S*|*=* |*Â*|from the input subnetwork family 𝒮 and with the largest total vertex score Σ_*v*∈*S*_ *X*_*v*_. By default, NetMix2 uses the *propagation family 𝒮* = ℳ_*δ,p*_. In this case, NetMix2 constructs an additional graph (the similarity threshold graph) based on vertex similarities quantified by Personalized PageRank from the input graph. The choice of subnetwork family 𝒮 for NetMix2 is flexible and can be generalized to other families defined by linear or quadratic constraints including the *connected family 𝒞*_*G*_, *edge-dense family ℰ*_*G,p*_, and *cut family 𝒯*_*G,ρ*_.

NetMix2 consists of two steps. As in our previous method NetMix [60], the first step is to estimate the number |*Â*| of vertices in the altered subnetwork *A*. NetMix estimated |*Â*| by fitting the vertex scores {*X*_*v*_}_*v*∈*V*_ to a Gaussian mixture model (GMM), under strict parametric assumptions on the altered distribution 𝒟_a_ = *N*(*μ*, 1) and background distribution 𝒟_b_ = *N* (0, 1). However, not all vertex score distributions are well-fit by normal distributions of this form. Thus, in NetMix2, we extend NetMix by using local false discovery rate (local FDR) methods [69, 71, 70] to estimate *α*, as local FDR methods make mild assumptions on the forms of the distributions 𝒟_a_, 𝒟_b_ (see the Supplementary Notes for further details).

The second step of NetMix2 is to compute the subnetwork *S* ∈ 𝒮 with size |*S*| = | *Â*| and largest total vertex score *X*_*v*_:

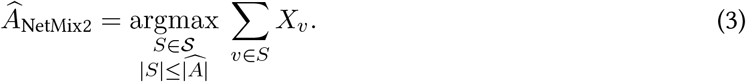

(3) can be computed using an integer linear program or integer quadratic program solver (e.g., Gurobi [83]) for a number of subnetwork families, including the edge-dense family *ℰ*_*G,p*_, the cut family 𝒯_*G,ρ*_, the connected family 𝒞_*G*_, and the propagation family ℳ_*δ,p*_. We note that for the propagation family ℳ_*δ,p*_, the run-time for solving (3) depends on both the number |*E*_*δ*_| of edges in the similarity threshold graph *G*_*δ*_ and the density *p*.

Note that (3) involves maximizing the sum Σ _*v*∈*S*_ *X*_*v*_ of the vertex scores *X*_*v*_, while the objective in the NetMix optimization problem [60] is the sum Σ _*v*∈*S*_ *r*_*v*_ of the vertex responsibilities *r*_*v*_ = *P*(*v* ∈ *A* | *X*_*v*_). In practice, we observe that maximizing the sum of the vertex scores *X*_*v*_ yields slightly better performance than that of the responsibilities *r*_*v*_.

### 2.4 Scores-only and network-only baselines

When evaluating any algorithm for the identification of altered subnetworks, we argue that it is essential to compare against two baselines: a *“scores-only”* baseline that only uses the vertex scores *X*_*v*_, and a *“network-only”* baseline that only uses the interaction network *G*. These two baselines quantify whether the altered subnetwork algorithm is outperforming simpler approaches that do not integrate vertex scores with a network; moreover, these baselines should be evaluated on each dataset and match as closely as possible the inputs to the altered subnetwork problem. A scores-only baseline is straightforward: we rank the vertices *v* by their vertex scores *X*_*v*_. Because this baseline outputs a ranked list of all vertices in the graph, we threshold the ranking when evaluating against other altered subnetwork algorithms by taking the *k* most highly ranked vertices for some integer *k*.

Defining a network-only baseline is a more subtle issue, and was discussed in two recent papers [68, 67]. Levi et al. [68] benchmarks altered subnetwork algorithms on randomly permuted vertex scores 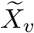 while keeping the network *G* fixed. The authors find that many existing methods output similar altered subnetworks (in terms of GO enrichment) on their permuted data, which suggests that these methods are utilizing the network *G* more than the vertex scores *X*_*v*_. Lazareva et al. [67] benchmarks altered subnetwork algorithms on randomly permuted networks with the same degree distribution as *G* while keeping the vertex scores *X*_*v*_ fixed. The authors find that many existing algorithms output similar altered subnetworks on permuted networks, indicating a degree bias in these methods. We propose a more direct network-only baseline: we rank vertices *v* by their network centrality score *N*(*v*) for a network centrality measure *N* that is derived from the topological constraints used by the altered subnetwork algorithm. For example, for an algorithm that relies on the connected subfamily, we propose that degree centrality *N*(*v*) = *d*_*v*_ is an appropriate measure, as in [67]. However, for network propagation algorithms that use random walk with restart, we claim that the PageRank centrality *N*(*v*) = (*M*_PPR_ · **1**)_*v*_, where **1** ∈ ℝ^*n*^ is an all-ones vector, is the more appropriate network-only baseline. This is because compared to degree centrality, PageRank centrality better captures how network propagation methods use the interaction network *G*.

## 3 Results

We evaluated NetMix2 on simulated data and on real datasets including somatic mutations in cancer and genome-wide association studies (GWAS) from several diseases. Unless indicated otherwise, we ran NetMix2 with the propagation family ℳ_*δ,p*_ using the personalized PageRank matrix *M*_PPR_ with restart probability *r* = 0.4. We solved the integer program in (3) using the Gurobi optimizer [83]. We ran Gurobi for up to 24 hours, which typically results in a near-optimal solution for the protein-protein interaction networks *G* that we used. For all ranking methods (e.g., network propagation, scores-only, and network-only baselines), we estimated the altered subnetwork *Â* as the |*Â*_NetMix2_ |highest ranked vertices, where *Â*_NetMix2_ is the output of NetMix2.

### 3.1 Simulated data

We compare NetMix2, network propagation, a scores-only baseline, and a network-only baseline on simulated instances of the Altered Subnetwork Problem with various subnetwork families 𝒮 derived from the HINT+HI protein interaction network *G* = (*V, E*) [23, 60]. This network contains 15,074 vertices and around 170,000 edges. The edges *E* are a union of the protein-protein and protein-complex interactions between proteins from the HINT network [84] and the protein-protein interactions from the HI network [85]. For each instance, we randomly selected a subnetwork *A* ∈ 𝒮 of size |*A*| = 0.01*n* and drew a sample **X** ∼ ASD _𝒮_ (*A, 𝒟*_*a*_, *𝒟*_*b*_) with altered distribution *D*_a_ = *N* (1.8, 1.2) and background *D*_*b*_ = *N* (0.1, 0.9). Note that we use slightly different altered and background distributions compared to the normally distributed ASD, so as to reflect the systemic errors in measurement often found in real data [69, 71, 70]. We implant altered subnetworks *A* from four different subnetwork families 𝒮: the connected family 𝒮= 𝒞 _*G*_, the edge density family 𝒮= *ℰ*_*G,p*_ with density *p* = 0.15, the cut family 𝒮= 𝒯 _*G,ρ*_ with cut-size *ρ* = 6, and the propagation family 𝒮= ℳ_*δ,p*_ with *δ* chosen so that *G*_*δ*_ has 200, 000 edges and with edge-density *p* = 0.3 in *G*_*δ*_. The edge-density (resp. cut-size) parameters are chosen to be at least 3 standard deviations above (resp. below) the average edge-density (resp. cut-size) for a subnetwork *A* of size |*A*| = 0.01*n*.

We ran each method on vertex scores **X**, with NetMix2 also having the subnetwork family 𝒮 as input, to obtain an estimate *Â* of the altered subnetwork *A*. We found that NetMix2 outperformed the other three methods — network propagation, scores-only, and network-only — for altered subnetworks *A* ∈ 𝒮 from all four families 𝒮 (Figure 2). However, the advantage of NetMix2 over the other methods depended on the subnetwork family 𝒮. For example, NetMix2 only slightly outperformed the scores-only baseline for the connected family 𝒞_*G*_ and cut family 𝒯_*G,ρ*_, and only moderately outperformed the scores-only baseline for the edge-dense family *ℰ*_*G,p*_, suggesting that these families impose relatively weak constraints on the altered subnetwork *A*, as discussed for the connected family in [60]. On the other hand, NetMix2 substantially outperformed the scores-only baseline for the propagation family ℳ_*δ,p*_, demonstrating that the propagation family ℳ_*δ,p*_ provides strong topological constraints on the altered subnetwork.

**Figure 2:**
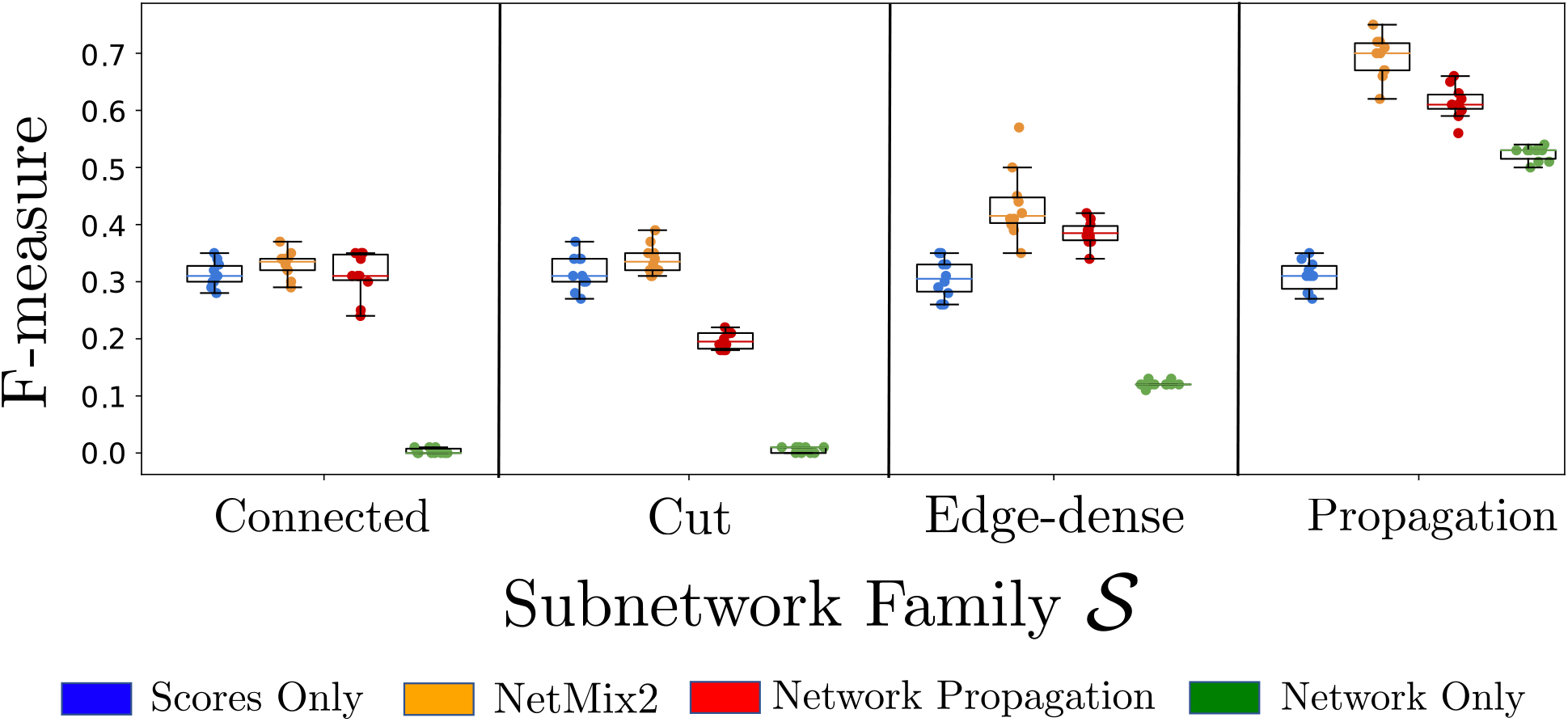
Comparison between NetMix2, network propagation, a scores-only baseline, and the PageRank centrality network-only baseline in identifying altered subnetworks from different subnetwork families 𝒮 implanted in the HINT+HI interaction network.

We also observed (Figure 2) that the performance of network propagation depends on the subnetwork family 𝒮. In particular, network propagation had similar performance to the scores-only baseline for the connected family 𝒞_*G*_ and edge-dense family *ℰ*_*G,p*_ and worse performance for the cut family 𝒯_*G,ρ*_, suggesting that network propagation is not well-suited to identifying altered subnetworks *A* from these subnetwork families. On the other hand, network propagation had a substantial gain over the scores-only baseline for the propagation family ℳ_*δ,p*_. This demonstrates that the propagation family contains subnetworks whose vertices are likely to be highly ranked by network propagation. Interestingly, we observe that for altered subnetworks *A* from the propagation family ℳ_*δ,p*_, the network-only baseline also outperforms the scores-only baseline. This is most likely due to the close relationship between the propagation family ℳ_*δ,p*_ and the network-only baseline: the network-only baseline ranks vertices by their PageRank centrality while the propagation family ℳ_*δ,p*_ is derived from the personalized PageRank matrix. Interestingly, ranking vertices by their degrees – as suggested in [67] – performed much worse than PageRank centrality (*F* -measure ≈ 0.05, not shown in Figure 2) for altered subnetworks *A* from the propagation family ℳ_*δ,p*_, demonstrating the importance of using a network-only baseline that models the same topological properties as the network method.

### 3.2 Somatic mutations in cancer

Next, we compared the performance of NetMix2 against several other methods for identification of altered subnetworks [8, 60, 86, 64] on the task of identifying cancer driver genes. For each vertex (gene) *v*, the vertex score *X*_*v*_ is a z-score computed from p-values from MutSig2CV [87], a statistical method that predicts cancer driver genes based on the frequency that the gene is mutated in a cohort of cancer patients. We obtained these scores for 10,437 samples across 33 cancer types from the TCGA PanCanAtlas project [88]. We also compared to three ranking methods: network propagation, scores-only, and network-only (PageRank centrality). We ran each method using the STRING protein-protein interaction network [89] and evaluated the performance by computing the overlap between genes in their reported subnetworks and reference lists of cancer driver genes from the COSMIC Cancer Gene Census (CGC) [90], OncoKB [91], and TCGA [88]. Further details on datasets and procedures for running each method are included in the Supplement.

We found that NetMix2 using the propagation family outperformed other methods in F-measure for all three reference gene sets (Table 1). In addition, comparing NetMix2 using the propagation family and NetMix2 using the connected family (the second best method) shows that the altered subnetwork found using the propagation family contains several genes that are not found by using the connected family (Figure S1). For example, NetMix2 using the propagation family identifies 9 CGC driver genes that are not found by using the connected family including *PDGFRA*, an oncogene whose gain-of-function mutations promote cancer growth [92] and *NCOR2*, a well-known tumor suppressor implicated in breast and prostate cancers [93]; none of these genes are found by the baseline methods (Figure S2). We also found that NetMix2 outperforms the altered subnetwork methods with different parameter settings including Heinz with FDR = 0.005 as well as the network-only baseline with degree centrality (Table S2). In particular, ranking vertices by degree had a substantially lower F-measure compared to ranking vertices by PageRank centrality – again demonstrating the importance of an appropriately chosen network-only baseline. We also observe that many network approaches, including NetSig and Hierarchical HotNet, had lower F-measure than the scores-only baseline. While it is possible we were not using the optimal parameters for these methods, our results suggest that these methods were over-utilizing the network compared to the vertex scores.

**Table 1:**
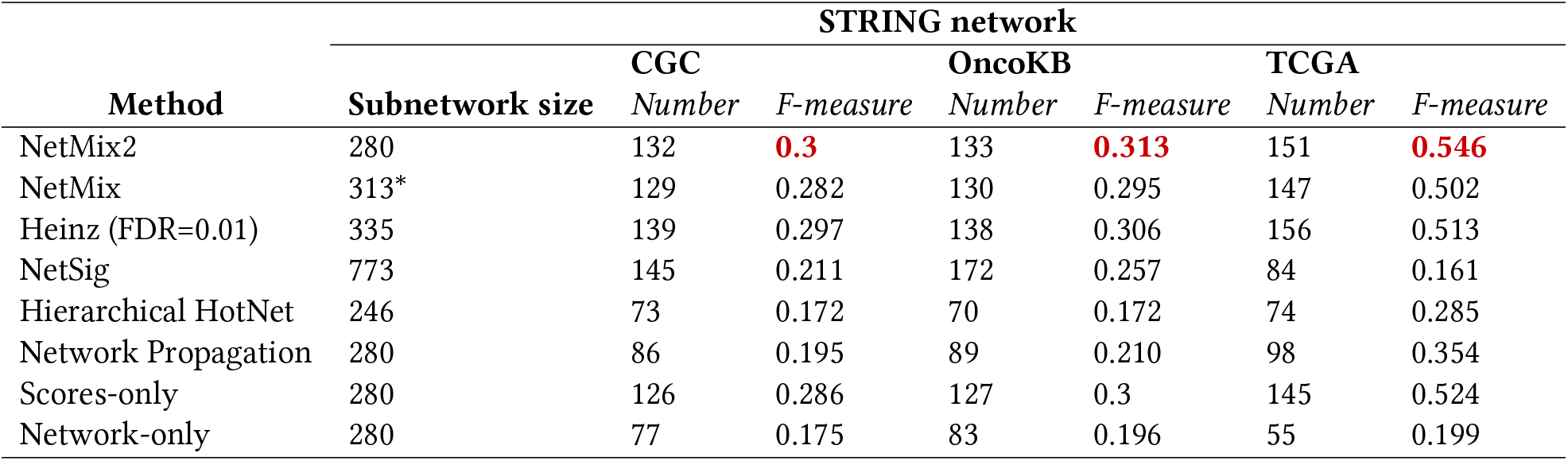
Results of altered subnetwork identification methods using MutSig2CV cancer driver gene *p*-values from TCGA tumor samples. Subnetworks are evaluated using reference sets of cancer genes from CGC, OncoKB, and TCGA. The best scores are colored in bold red. ^*^*GMM from NetMix overestimated the size of the subnetwork, thus we excluded genes with outlier scores as described in [60]*.

| Method | Subnetwork size | STRING network |  |  |  |  |  |
| --- | --- | --- | --- | --- | --- | --- | --- |
|  |  | CGC |  | OncoKB |  | TCGA |  |
|  |  | Number | F-measure | Number | F-measure | Number | F-measure |
| NetMix2 | 280 | 132 | **0.3** | 133 | **0.313** | 151 | **0.546** |
| NetMix | 313* | 129 | 0.282 | 130 | 0.295 | 147 | 0.502 |
| Heinz (FDR=0.01) | 335 | 139 | 0.297 | 138 | 0.306 | 156 | 0.513 |
| NetSig | 773 | 145 | 0.211 | 172 | 0.257 | 84 | 0.161 |
| Hierarchical HotNet | 246 | 73 | 0.172 | 70 | 0.172 | 74 | 0.285 |
| Network Propagation | 280 | 86 | 0.195 | 89 | 0.210 | 98 | 0.354 |
| Scores-only | 280 | 126 | 0.286 | 127 | 0.3 | 145 | 0.524 |
| Network-only | 280 | 77 | 0.175 | 83 | 0.196 | 55 | 0.199 |

### 3.3 Genome Wide Association Studies (GWAS)

We next applied NetMix2 to data from genome-wide association studies (GWAS), another application where network approaches are often used to prioritize germline variants that are associated with biological traits [37, 96]. We first analyzed GWAS data for eight different disease traits from [94]. The original study [94] introduced the NAGA method (also in companion paper [97]) which outputs a *ranked list* of genes by applying network propagation to gene scores *X*_*v*_ obtained by selecting the single nucleotide polymorphism (SNP) with minimum *p*-value in the neighborhood of each gene *v*. The genes are then ranked by their propagated scores *Y*_*v*_, and these rankings are evaluated using reference lists of disease genes from the DisGeNET database [95].

As a preliminary analysis, we first compared NAGA (network propagation) against our scores-only and network-only (PageRank centrality) baselines. The original study [94] claimed that network propagation outperforms the scores-only baseline (as well as existing network ranking methods for GWAS) in recovering reference genes for all eight diseases. They demonstrated their claim by comparing network propagation against other ranking methods (rather than as altered subnetwork methods) using the area under the receiver operating characteristic curve (AUROC) metric. However, when the reference gene list is small (which is typically the case for GWAS data), the AUROC of a ranking algorithm is not necessarily representative of its performance; two network ranking algorithms can have similar AUROC even if they have noticeably different performance [98]. Therefore, we re-evaluated the network ranking approaches from [94] for prioritizing disease genes using area under the precision-recall curve (AUPRC), a metric which is reported to be a more appropriate metric for assessing ranking algorithms compared to AUROC when the reference sets are small [98]. We focused our comparison on network propagation (using the same parameters as NAGA), the scores-only baseline, and the network-only baseline using PageRank centrality. We used the PCNet interaction network *G* [18], the same network used in the original evaluation [94].

We found that – contrary to the claims of [94] – network propagation does not always outperform the scores-only baseline (Figure 3). Furthermore, based on the additional comparison to the network-only baseline, we identified three distinct groups of GWAS datasets (Figure 3). In the first group are three diseases (schizophrenia, hypertension, type 1 diabetes) where the network-only AUPRC was comparable (less than 10% difference) to the network propagation AUPRC. In other words, the first group consists of datasets where the vertex scores *X*_*v*_ did not seem to add much value compared to using only the network *G* in identifying reference genes. Interestingly, this group includes schizophrenia, a disease which [94, 66] specifically highlight as a case where network propagation outperformed the scores-only baseline in AUROC. The second group consists of two diseases (Crohn’s disease, coronary artery disease) where the scores-only AUPRC was comparable to (or larger than) the network propagation AUPRC. This group consists of diseases where the network *G* seemingly did not add much value compared to using only the vertex scores *X*_*v*_ in identifying reference genes. Finally, the third group consists of three diseases (Rheumatoid arthritis, bipolar disorder, type 2 diabetes) where the network propagation AUPRC is noticeably larger (at least 10%) than the AUPRC for both scores-only and network-only. This group consists of diseases where using both the network *G* and scores *X*_*v*_ was better at identifying reference genes than either alone, and is the group of diseases where one would expect altered subnetwork approaches to perform well. We also note that for all eight diseases, ranking vertices by their degree (not shown in Figure 3) had lower AUPRC than ranking vertices by their PageRank centrality, again illustrating the benefit of comparing to appropriate network-only baselines.

**Figure 3:**
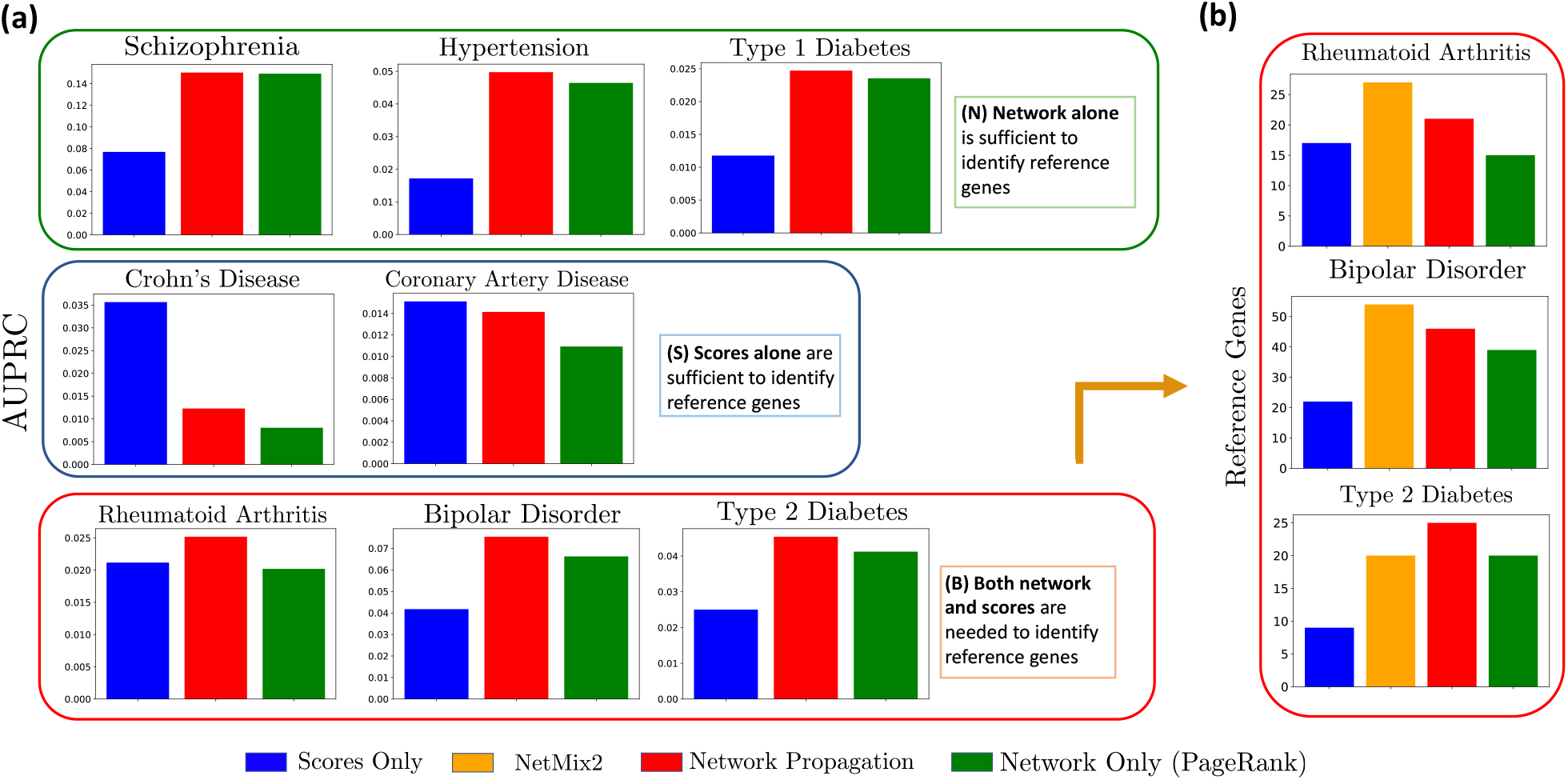
**(a)** Comparison of three network ranking methods — network propagation, scores-only, and network-only (PageRank centrality) — on GWAS data from [94]. Network ranking methods are evaluated by their AUPRC using reference lists of disease genes [95]. **(b)** Comparison of four altered subnetwork identification methods — NetMix2, network propagation, scores-only, and network-only (PageRank centrality) — for diseases from (a) where network propagation has at least 10% larger AUPRC compared to the scores-only and network-only baselines. Methods are evaluated according to the number of reference genes in the estimated altered subnetwork.

Next we compared NetMix2 against network propagation and the two baselines for the three diseases in the latter group: Rheumatoid arthritis, bipolar disorder, and type 2 diabetes (Figure 3b). We found that for two out of three diseases (rheumatoid arthritis and bipolar disease) NetMix2 identified noticeably more reference genes than network propagation. These results are consistent with the simulations in Figure 2, and demonstrate that NetMix2 using the propagation family outperforms network propagation for the diseases where *both* the scores and the network are important for identifying reference disease genes.

One empirical observation from comparing the subnetworks identified by different approaches is that the NetMix2 subnetwork was similar to the scores-only subnetwork, while the network propagation subnetwork was more similar to the network-only subnetwork (Figure S3). As a result, we expect that the quality of the gene scores *X*_*v*_ computed from the GWAS summary statistics of individual SNPs plays an important role for the performance of NetMix2 as well as the scores-only baseline. Recent studies [99, 100] report that the minSNP method – which finds the most significant SNP in surrounding regions of each gene and was used by [94] to compute the gene scores in this experiment – introduces bias towards longer genes and does not account for linkage disequilibrium, a well known confounding factor in GWAS summary statistics [101].

To address this issue, we examined another set of GWAS [68] where gene scores are computed using the *Pascal* method [99]. Briefly, Pascal aggregates SNP *p*-values from GWAS into gene scores while correcting for confounders including linkage disequilibrium and gene length. We applied NetMix2 to Pascal scores of six diseases from the GWAS data in [68]. We used the STRING interaction network [89], the same network used in [68], and compared against three ranking methods — network propagation, the scores-only baseline, and the network-only (PageRank centrality) baseline — as well as DOMINO, the altered subnetwork identification method presented in [68]. Similar to the previous experiment, we evaluated the methods by computing the overlap between subnetworks identified by each method and DisGeNET reference disease gene sets [95].

We found that the Pascal gene scores alone represent a strong signal for recovering disease reference genes as the scores-only baseline outperformed other methods in four of the six diseases, namely atrial fibrillation, coronary artery disease, age-related macular degeneration, and Crohn’s disease (Figure 4). Consistent with the results on the previous GWAS experiment, we also found that the NetMix2 subnetwork was more similar to the scores-only subnetwork, while the network propagation subnetwork was more similar to the network-only subnetwork (Figure S4). Thus, it is not surprising to see that NetMix2 performed nearly as well as the scores-only baseline in these four diseases. The strong performance of scores-only baseline using Pascal scores demonstrates that the quality of gene scores is an important component of the performance of altered subnetwork methods.

**Figure 4:**
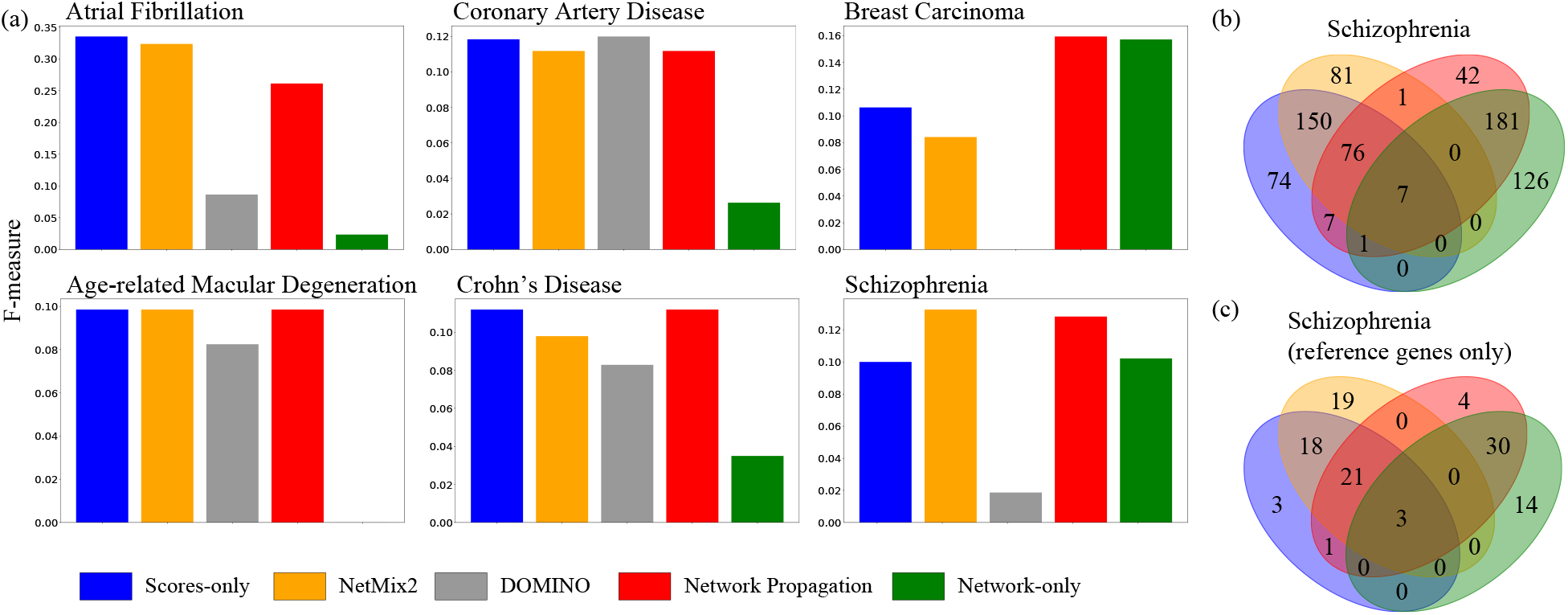
**(a)** Comparison of NetMix2, DOMINO, network propagation, and the scores-only and network-only (PageRank centrality) baseline methods on GWAS data from [68] using the STRING interaction network [89] **(b)** Overlap in genes in altered subnetworks in schizophrenia identified by scores-only, NetMix2, network propagation, and network-only. **(c)** Schizophrenia reference genes in the altered subnetworks from (b).

We next evaluated DOMINO against the scores-only and network-only baselines – comparisons that were missing from the DOMINO publication [68] – and against NetMix2. We found that DOMINO performed worse than the baseline methods as well as NetMix2 in five of the six diseases, with coronary artery disease being the lone exception (Figure 4). On the other hand, NetMix2 was comparable to the best performing methods in four diseases and also higher than all other methods in schizophrenia (Figure 4a). For schizophrenia, the altered subnetwork identified by NetMix2 contained 81 genes that were not found by the ranking methods (Figure 4b). These 81 genes are significantly enriched (*p*-value = 3.2e-05, fold enrichment = 2.54; hypergeometric test) for the reference genes for schizophrenia from the DisGeNet database (Figure 4c). Furthermore, the altered subnetworks identified by NetMix2 for schizophrenia as well as breast carcinoma are significantly enriched (schizophrenia: *p*-value = 2.04e-11, fold enrichment = 16.1; hypergeometric test, and breast carcinoma: *p*-value = 1.56e-22, fold enrichment = 6.38; hypergeometric test) for genes whose expression is significantly associated with the corresponding diseases according to recent studies of expression quantitative trait loci (eQTL) and GWAS data [102, 103]. Lastly, we note that the results were qualitatively similar when we used the size of the subnetwork identified by DOMINO (instead of the subnetwork found by NetMix2) to threshold the ranking methods (Figure S5).

Taken together, these results demonstrate the importance of evaluating methods for identifying altered subnetwork against the scores-only or the network-only baseline methods, both to gauge the potential bias from either source of information and to comprehensively assess the performance of a network method.

## 4 Discussion

We introduced NetMix2, an algorithm that unifies the network propagation and altered subnetwork approaches to analyze biological data using interaction networks. NetMix2 is inspired by network propagation, a standard approach for solving the *network ranking* problem, and attempts to bridge the gap between two paradigms for using networks in the analysis of high-throughput genomic data — *network ranking* and the identification of *altered subnetworks* — in a principled way by explicitly deriving a new family of subnetworks called the *propagation family* that approximates the altered subnetworks found by network propagation methods. We showed that NetMix2 is effective in finding disease-associated genes using somatic mutation data in cancer and GWAS data from multiple diseases.

At the same time, our evaluation revealed that simple baseline methods that use either only the vertex scores or only the interaction network sometimes perform surprisingly well, often outperforming more sophisticated network methods. While publications describing new network methods typically benchmark against other network methods, they are wildly inconsistent in benchmarking against scores-only and network-only baselines. It is rare to see a paper that benchmarks against both baselines. Moreover, the network-only baseline should be calibrated to use the same network information as the method under evaluation; e.g., PageRank centrality is a more appropriate benchmark for network propagation methods than vertex degree.

While NetMix2 outperformed every other method we tested on cancer data, the performance of NetMix2 – and other network methods – was underwhelming in general on the GWAS data, with modest improvement over baseline methods even in the diseases where NetMix2 worked well. There are several possible reasons for this discrepancy. First, the diseases in the GWAS data that we analyzed may have high genetic complexity. Indeed, multiple studies have suggested that the signal from GWAS data for complex traits are more widely dispersed throughout the genome (including non-coding genomic regions) than previously predicted, resulting in very small effect sizes of individual entities (SNPs/genes). An extreme example is the omnigenic model [104] which posits that nearly all genes have functional relevance to the GWAS trait. Such complexity coupled with various challenges in interpreting the GWAS data, e.g., linkage disequilibrium, could result in a high number of false positives — genes with significant *p*-values that are not related to the trait — or a gene score distribution with an unusual shape that is not well fit by the semi-parametric models in local FDR methods. This could result in an inaccurate estimation of the size of the altered subnetwork which in turn would be detrimental to the performance of NetMix2 as well as the ranking methods that select the same number of genes as NetMix2, since the relative difference in size between the estimated altered subnetwork and the reference gene set imposes an inherent upper-bound on the performance of an algorithm.

There are several directions for future work. The first direction is to extend NetMix2 to identify *multiple* altered subnetworks simultaneously. This can be done by running NetMix2 iteratively, or by modifying the integer program to output multiple solutions. However, solving the corresponding model selection procedure to choose the number and sizes of altered subnetworks without overfitting is a difficult problem. A second direction is to extend NetMix2 with an appropriate permutation test to evaluate the statistical significance of the altered subnetwork(s). Finally, while we evaluated several network methods and simple baselines, there are numerous other network methods that could be included in these benchmarks. However, there are few gold standards to perform such a comprehensive evaluation as the reference disease gene sets remain relatively limited and potentially biased by their source. Thus, a useful extension would be deriving a reliable evaluation scheme for network methods that accounts for various sources of bias including the ascertainment bias in current interaction networks and disease gene sets.

## 4.1 Acknowledgements

The authors would like to thank Jasper C. H. Lee and Christopher Musco for helpful discussions, as well as Matthew Myers and Palash Sashittal for reviewing early versions of the manuscript. U.C. is supported by NSF GRFP DGE 2039656. B.J.R. is supported by grant U24CA264027 from the National Cancer Institute (NCI).

## Supplementary Notes

### Proof of Proposition 1

#### Lemma 2.

*Let* 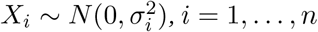 *be independently distributed Gaussians with bounded variance* 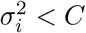. *Then*

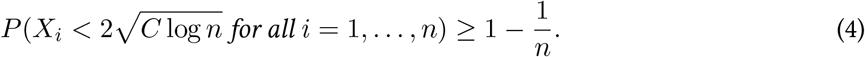

*Proof*. For fixed *i* ∈ [*n*], we have

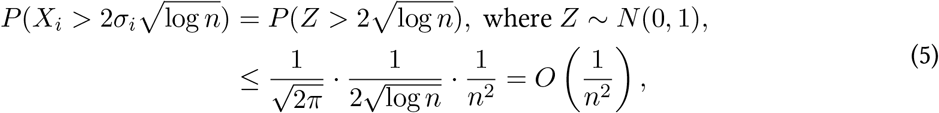

where in the last inequality we use the standard bound. 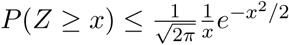 Thus,

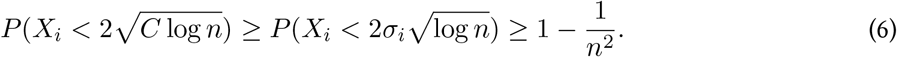

By a union bound, it follows that

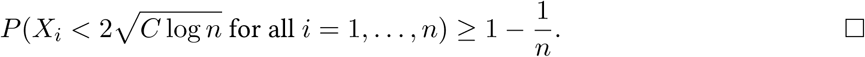

#### Proposition 1.

*Let M* ∈ [0, 1]^*n*×*n*^ *be a matrix, and define r* = min_*v*∈*V*_ *M*_*v,v*_, *c* = max_*v* ∉*V*_ Σ_*w*∈*A*_ *M*_*v,w*_, *and d* = max_*v,w*∈*V*_ Σ_*u*∈*V*_ (*M*_*w,u*_−*M*_*v,u*_)^2^. *Let* 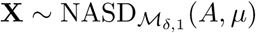 *where μ* ≥ 1 *and* 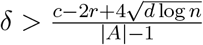. *Then with probability at least* 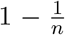, *the altered subnetwork A consists of the* |*A*| *vertices with the largest propagated scores Y*_*v*_.

*Proof*. First, we define the random variables {*Z*_*v*_}_*v*∈*V*_ as

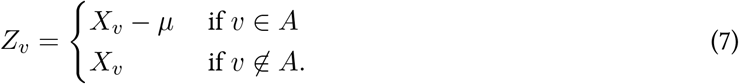

Note that *Z*_*v*_ ∼ *N* (0, 1) for all vertices *v* ∈ *V*.

Now, we are done if we prove that *Y*_*a*_ *> Y*_*b*_ for all *a* ∈ *A* and *b* ∉ *A* with probability at least 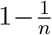. Recall that the propagated score *Y*_*v*_ is defined as *Y*_*v*_ = Σ_*w*∈*V*_ *M*_*v,w*_ *X*_*w*_. Using the definition of the propagated scores *Y*_*v*_, an equivalent claim is

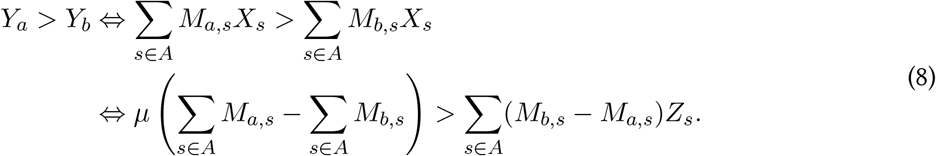

To prove (8), we start by lower bounding the LHS of (8). We have

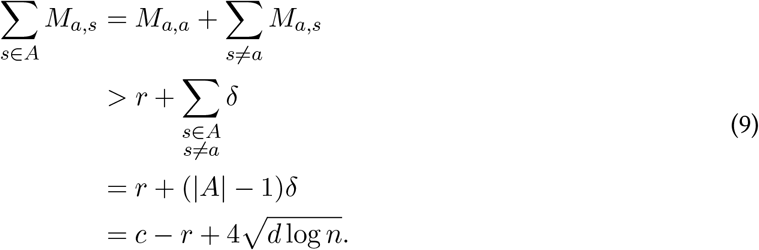

We also have

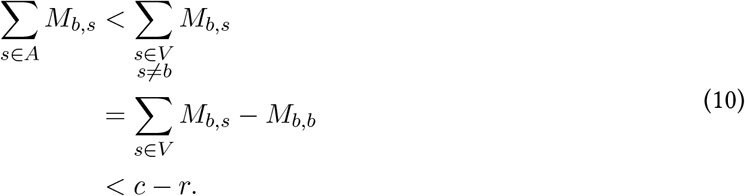

Thus, using that *μ* ≥ 1, we have

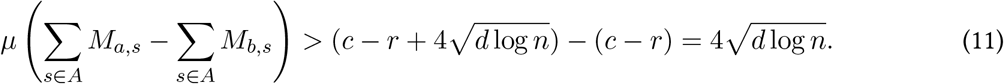

Next we upper bound the RHS of (8). We have that

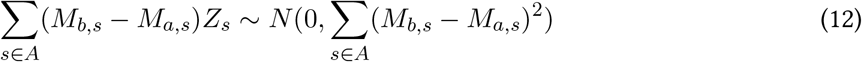

where Σ_*s*∈*A*_(*M*_*b,s*_ − *M*_*a,s*_)^2^ ≤ *d*. Thus, by Lemma 2, we have that

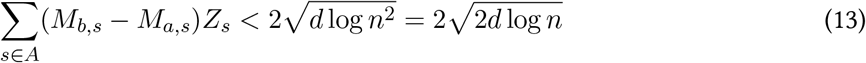

for all *a* ∈ *A, b* ∉ *A* with probability at least 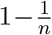. It follows that

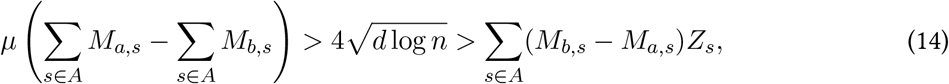

thus proving (8).□

### Estimating the size of altered subnetwork using local FDR

The first step of NetMix2 is to estimate the size of the altered subnetwork *A* using local FDR methods [69, 70, 71, 105]. While the original local FDR methods estimate the background and altered distributions of vertex scores, they assume that the altered distribution is *two-sided*. Therefore, we derive a heuristic approach using the results of local FDR methods to estimate the number of vertices in a *one-sided* altered distribution. Specifically, we first fit the vertex scores **X** = (*X*_*v*_)_*v*∈*V*_ using local FDR methods [69, 70, 71, 105] which yields (1) an estimate 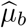 of the mean of the background distribution 𝒟_*b*_ and (2) estimates 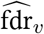 of fdr_*v*_ = *P*(*v* ∉ *A* | *X*_*v*_) for each vertex *v* (this quantity is known as the local FDR for vertex *v*). We then estimate the size |*A*| of the altered subnetwork *A* as the number of vertices *v* with score 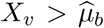 and estimated local FDR 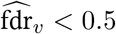, i.e.,

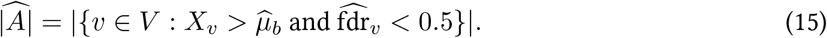

The first condition, 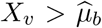, ensures that our estimate of |*A*| only counts vertices *v* with scores *X*_*v*_ that are larger than expected, consistent with our assumption that the altered distribution 𝒟_*a*_ is one-sided. The second condition, 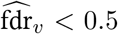, is equivalent to 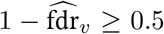, whose left-hand side is approximately 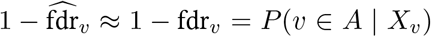. Thus, the second condition ensures that our estimate of |*A*| only counts vertices *v* with probability at least 0.5 of being in the altered subnetwork *A*.

### Data and Code Availability

NetMix2 code as well as a tutorial (Jupyter notebook) for running NetMix2 on example data are publicly available at https://github.com/raphael-group/netmix2.

### Parameters for running NetMix2

The input to NetMix2 are the network *G*, gene scores *X*_*v*_, and the subnetwork family *S*. For step 1 of NetMix2 (estimate the proportion 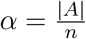 of vertices in the altered subnetwork *A*), we used the local FDR method with the default parameters. For step 2 (find the altered subnetwork *A*), the following additional parameters are required for the propagation family ℳ_*δ,p*_.

- *E*_*δ*_: the number of edges in the similarity threshold graph.
- *ρ*_*δ*_: the minimum edge density of the altered subnetwork.

In our analyses, we chose the values of each parameter for running NetMix2 to reflect the network properties of the input diseases. For all diseases, we chose the number |*E*_*δ*_| of edges in the similarity threshold graph *G*_*δ*_ to be roughly 40% of the number |*E*| of edges in the original protein interaction network *G*, for consistency in the similarity threshold graph *G*_*δ*_ across different diseases. However, we chose different values of the minimum edge density *ρ* of the altered subnetwork *A* for different diseases, depending on whether the gene scores or the network information contribute more to recovering disease-associated genes reported in reference databases. We use a larger value of *ρ* for diseases where the network-only baseline outperforms the scores-only baseline, i.e., diseases where the network information is more useful for finding disease-associated genes compared to the gene scores, and likewise we use a smaller value of *ρ*for diseases where the scores-only baseline outperforms the network-only baseline, i.e., diseases where the gene scores are more useful for finding disease-associated genes compared to the interaction network. The values of each parameter that we used for running NetMix2 are listed in Table S1. We ran NetMix2 for up to 24 hours unless stated otherwise.

### Data

#### Gene scores

For the analysis of somatic mutations in cancer, we used the MutSig2CV [106] driver *p*-values from the TCGA PanCanAtlas project [88]^2^. We also used the GWAS summary statistics (SNP *p*-values) from [94]^3^ and the Pascal gene scores from [68]^4^. For the GWAS data from [94], we mapped the SNP *p*-values to gene *p*-values using the NAGA software from [94] which selects the most significant SNP *p*-value within the 10 kilobases upstream and downstream regions for each gene. For all of our analyses, we used the *z*-scores as gene scores computed using *z* = Φ^−1^(*P*) where *P* is a *p*-value and Φ is the cumulative distribution function of the standard normal distribution.

#### Interaction networks

We used the following interaction networks for our analysis.

- HINT+HI [84, 85]
- STRING [89]
- PCNet [18]

We used the same procedures from [64] for constructing the HINT+HI network. For the STRING network, we selected interactions with a confidence score above 0.7. We also removed 6 and 1 vertices with the highest degrees from the HINT+HI and STRING networks, respectively, based on visual inspection of the degree distributions (Figure S6). For fair comparison with NAGA, we do not remove any vertices from PCNet.

#### Reference disease genes

For the analysis of somatic mutations in cancer, we used the lists of cancer driver genes from COSMIC Cancer Gene Census [90]^5^, OncoKB [91]^6^, and TCGA [88]. For the GWAS analyses, we used the “Curated” disease genes from the DisGeNet database [95]^7^.

### Evaluating network methods on somatic mutations in cancer

We ran 5 altered subnetwork identification methods — NetMix2, NetMix, Heinz, NetSig, and Hierarchical HotNet — and 4 ranking methods — network propagation, the scores-only baseline, the network-only baseline, and the degrees-only baseline on MutSig2CV *p*-values from the TCGA PanCanAtlas project using the STRING interaction network.

We ran NetMix2 using the propagation family and the connected family. The input parameters used for running NetMix2 using the propagation family are summarized in Table S1. Genes with extremely significant *p*-values skewed the fit of Gaussian mixture model in the original NetMix algorithm, resulting in an overestimate size of the altered subnetwork. Thus, we excluded 112 genes with *FDR <* 0.001 where *FDR* was computed using the Benjamini-Hochberg multiple testing correction [107]. We ran Heinz using two parameter settings, *FDR* = 0.01 and *FDR* = 0.005. We ran NetSig and Hierarchical HotNet with the default settings. For network propagation, we used Personalized PageRank with restart probability of 0.4. We used the same restart probability in other parts of this paper unless stated otherwise.

For all of the above methods, we restricted the input gene scores to the genes that are present in the network (except for network-only and degrees-only that do not use gene scores).

The results of each method are summarized in Tables 1 and S2. Figures S1 and S2 show comparisons between NetMix2 using the propagation family and the connected family and between NetMix2 and three ranking methods — network propagation, scores-only, and network-only.

### Evaluating network methods on GWAS data

For the eight diseases in GWAS data from NAGA [94], we conducted a preliminary analysis where we used the ranking methods — scores-only, NAGA (network propagation), and the network-only basline — to generate ranked lists of all genes in each disease using PCNet, the interaction network used in NAGA [108]. For the NAGA method, the original pipeline for propagating vertex scores in [94] used the same setting (Personalized PageRank with restart probability 0.4) as network propagation in our paper. Thus, we used our implementation of network propagation to compute the NAGA scores for ranking vertices.

Following the preliminary analysis, we ran NetMix2 and three ranking methods — NAGA (network propagation), scores-only baseline, and network-only baseline — to identify altered subnetworks in three of the eight diseases where network propagation performed well. We ran NetMix2 using the propagation family. The input parameters used for running NetMix2 on each GWAS data are summarized in Table S1.

We also analyzed six diseases in GWAS data from DOMINO [68]. For this dataset, we ran NetMix2, the same three ranking methods used for GWAS data from NAGA, and the DOMINO method (with default settings) using STRING, the interaction network used in DOMINO [68]. When applying the ranking methods on the DOMINO data, we used two different values— the size of the subnetwork identified by NetMix2 and by DOMINO — for selecting the top genes.

For all of the above methods, we restricted the input gene scores to the genes that are present in the network (except for network-only).

Comparisons of the subnetworks identified by each method for the GWAS data are summarized in Figures S3 and S4.

### Implementation and Software

We implement NetMix2 using Python 3. We use the Python implementation of locFDR R package from https://github.com/leekgroup/locfdr-python for step 1 of NetMix2. We use Gurobi software [83] for solving the integer linear/quadratic program in (3).

## Supplementary Figures

**Figure S1:**
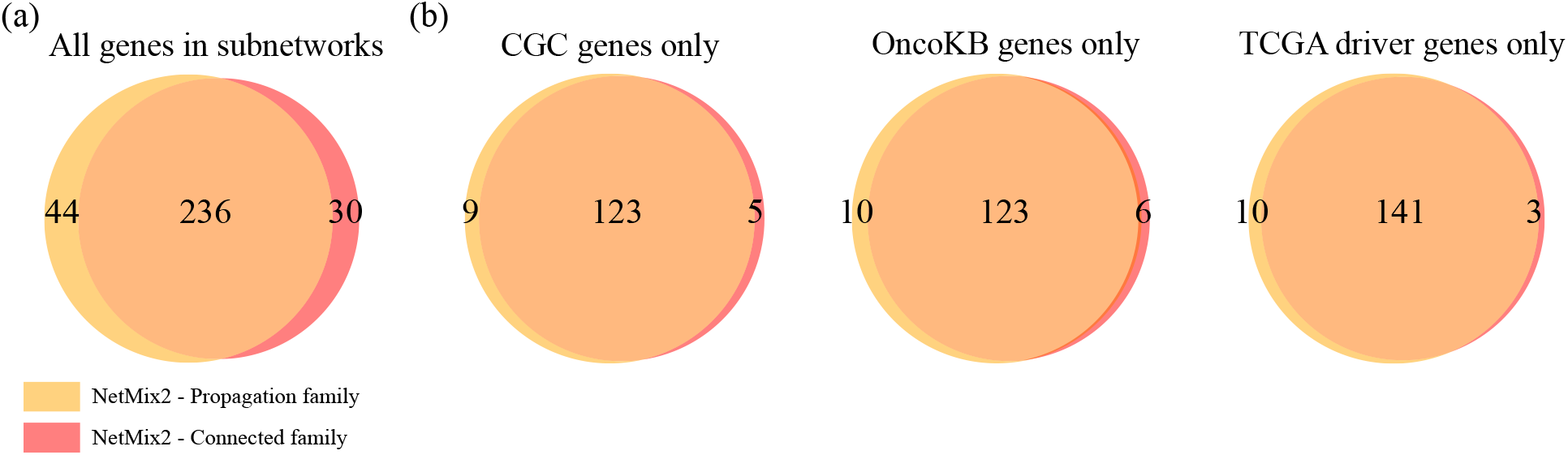
**(a)** Overlap between genes in the altered subnetworks identified by NetMix2 using similarity threshold family and by NetMix2 using connected family using MutSig2CV driver *p*-values from the TCGA PanCanAtlas project and the STRING network.. **(b)** Reference genes – COSMIC Cancer Gene Census genes, OncoKB cancer genes, and TCGA driver genes – in the altered subnetworks from (a).

**Figure S2:**
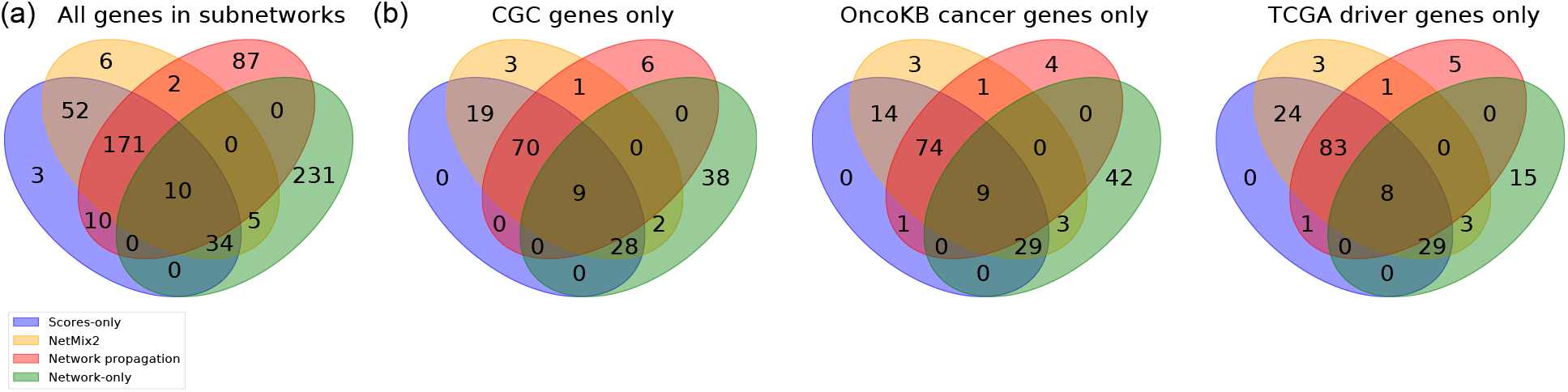
**(a)** Overlap in genes in altered subnetworks identified by the scores-only basline, NetMix2, network propagation, and the network-only basline using MutSig2CV driver *p*-values from the TCGA PanCanAtlas project and the STRING network. **(b)** Reference genes — CGC cancer genes, OncoKB cancer genes, and TCGA driver genes — in the altered subnetworks from (a).

**Figure S3:**
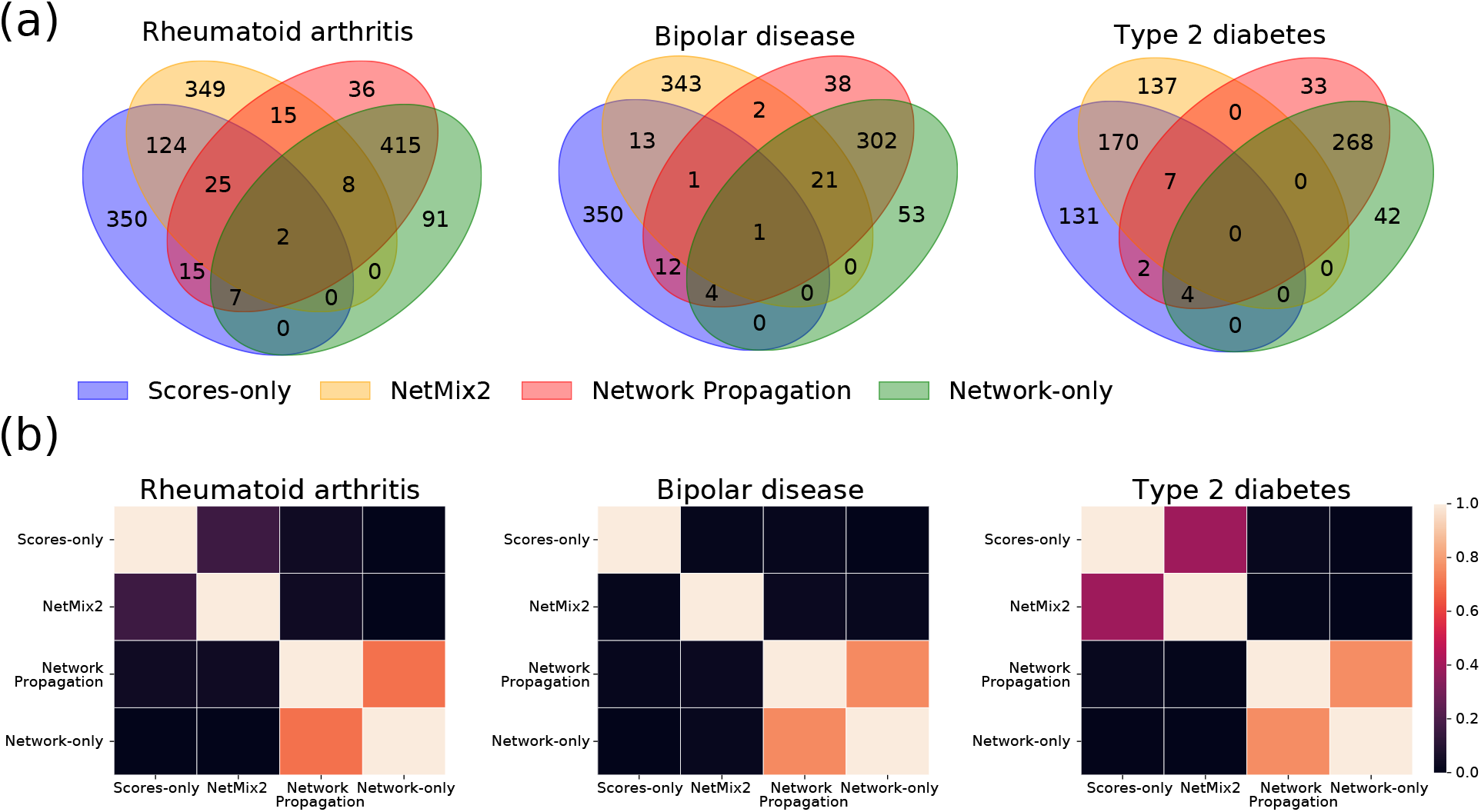
**(a)** Overlaps in genes in altered subnetworks identified by the scores-only basline, NetMix2, network propagation, and the network-only basline on the GWAS data from NAGA [94]. **(b)** Jaccard similarity matrix showing similarities between subnetworks from methods in (a). The subnetworks identified by NetMix2 are more similar to those identified by scores-only than network propagation and network-only.

**Figure S4:**
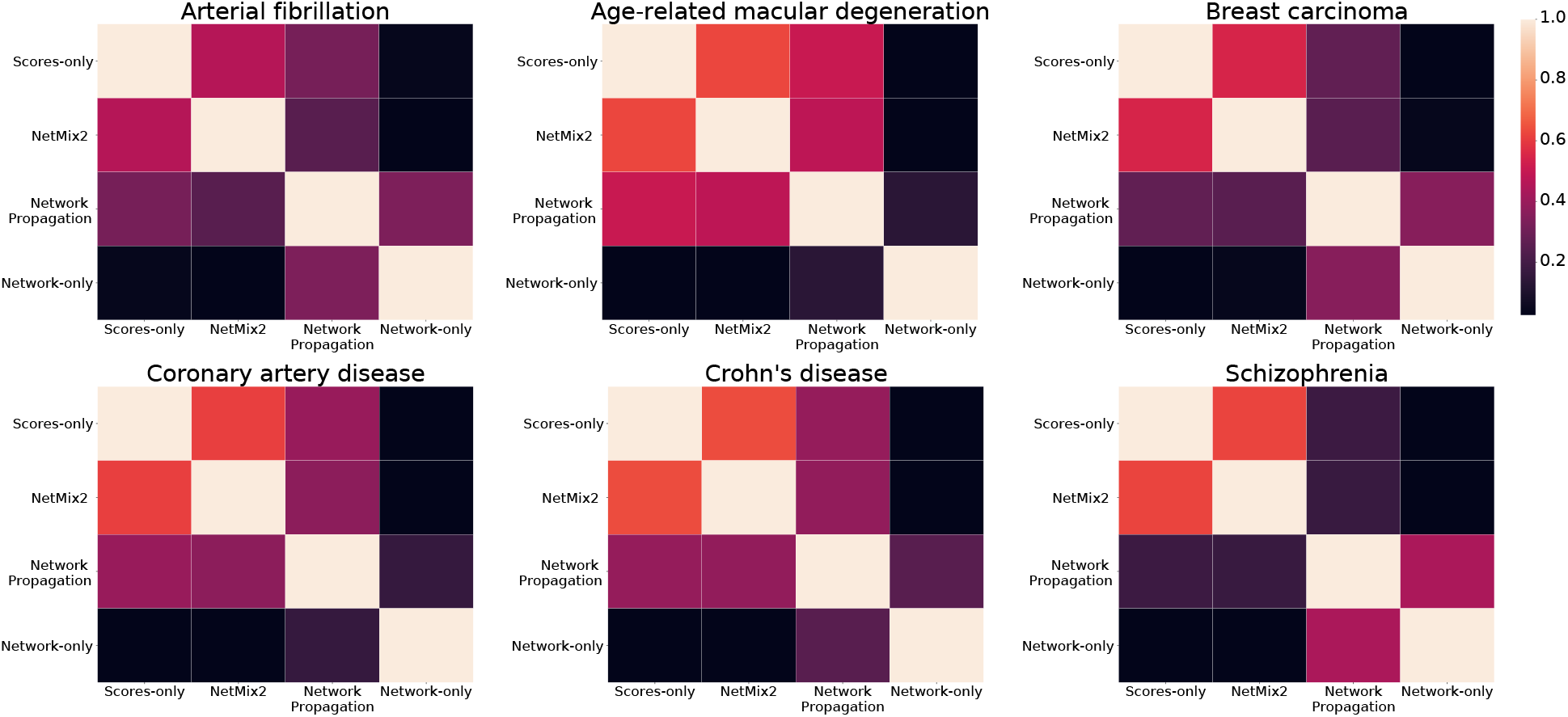
Jaccard similarity matrix showing similarities between subnetworks identified by different network methods on the GWAS from [68]. The subnetworks identified by NetMix2 are more similar to those identified by scores-only than network propagation and network-only.

**Figure S5:**
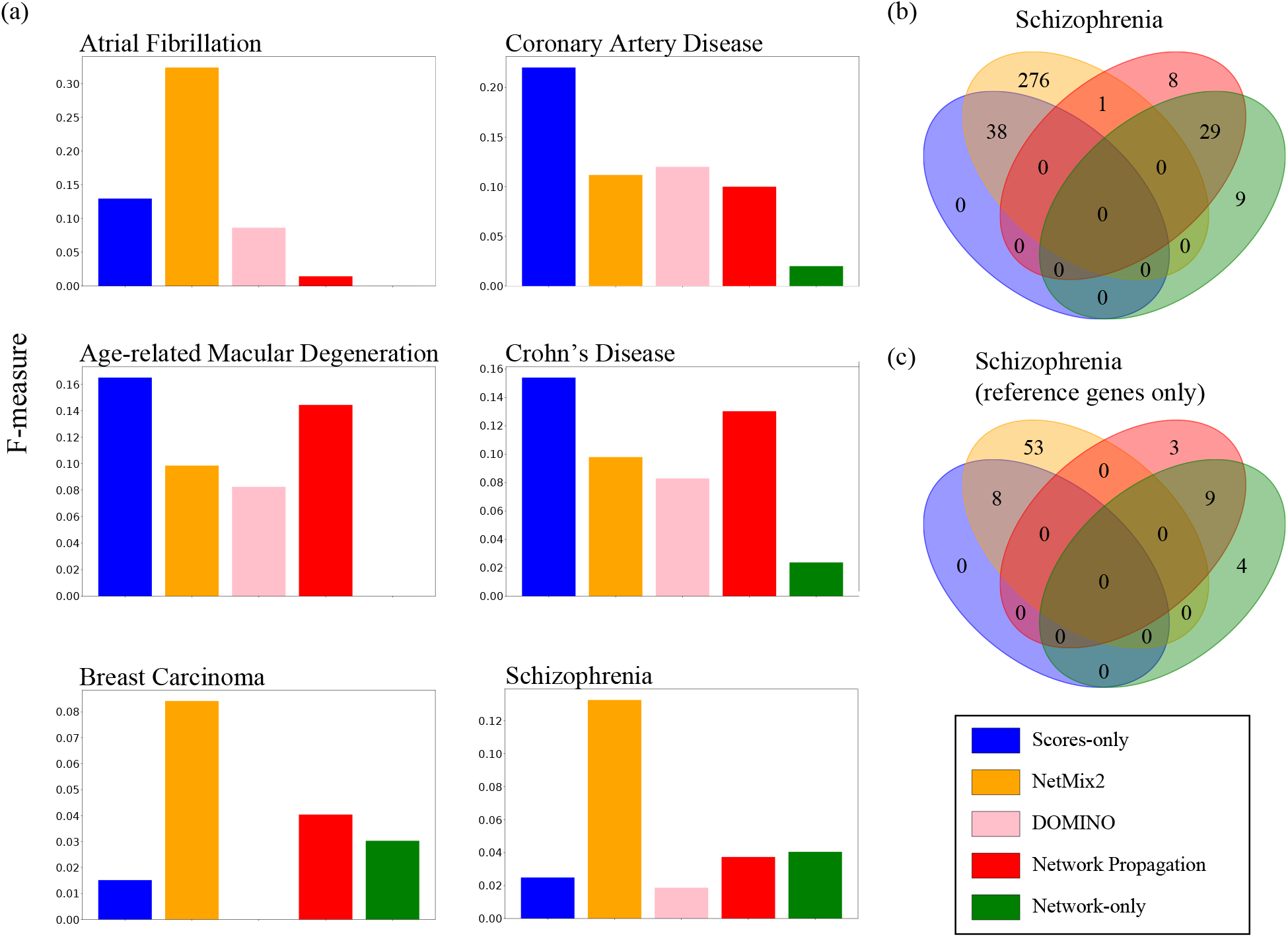
**(a)** Comparison of altered subnetwork methods — the scores-only basline, NetMix2, DOMINO [68], network propagation, and the network-only basline. For the ranking methods, top-*k* genes were selected where *k* = |*Â*_DOMINO_| is the size of the subnetwork identified by DOMINO |*Â*_DOMINO_|. **(b)** Overlap in genes in altered subnetworks in schizophrenia identified by scores-only, NetMix2, network propagation, and network-only. **(c)** Schizophrenia reference genes in the altered subnetworks from (a).

**Figure S6:**
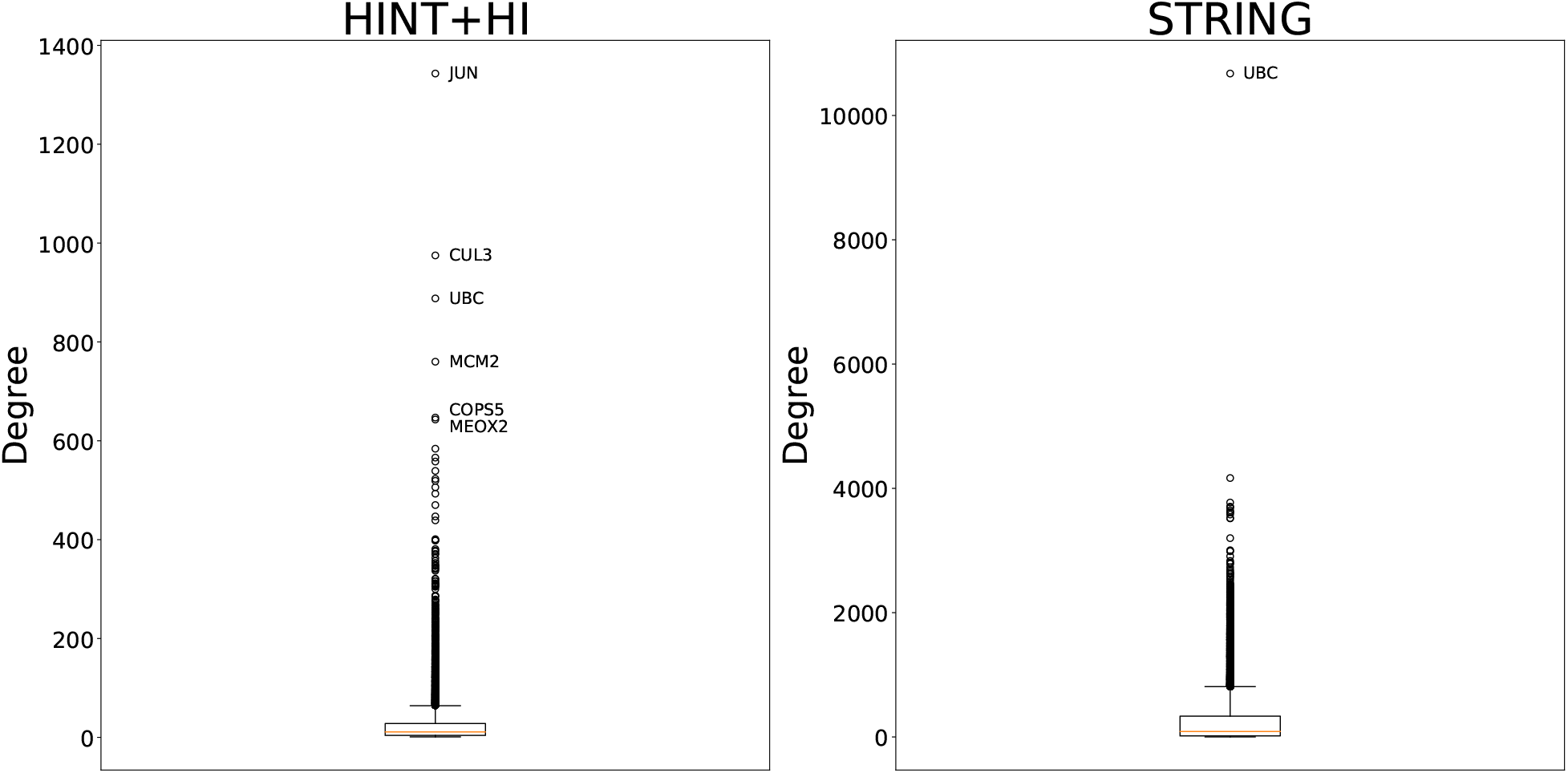
Distributions of the vertex degrees in HINT+HI network and STRING network. Removed vertices are labeled with gene names.

## Supplementary Tables

**Table S1:** List of datasets and parameters used in the reported results of NetMix2.

| | **Dataset** | **Network (size)** | $G_\delta$ size | **density $\rho$** | $\vert \hat{A}\vert $ |
| --- | --- | --- | --- | --- | --- |
| Pan-cancer somatic mutations | TCGA | STRING<br>(457516) | 150000 | 0.01 | 280 |
| Rheumatoid Arthritis | NAGA | PCNet<br>(2386616) | 1000000 | 0.05 | 523 |
| Bipolar Disease | NAGA | PCNet<br>(2419566) | 1000000 | 0.2 | 381 |
| Type 2 Diabetes | NAGA | PCNet<br>(2420208) | 1000000 | 0.2 | 314 |
| Atrial Fibrillation | DOMINO | STRING<br>(429272) | 175000 | 0.05 | 401 |
| Coronary Artery Disease | DOMINO | STRING<br>(401682) | 175000 | 0.05 | 258 |
| Age related Macular Degeneration | DOMINO | STRING<br>(301020) | 175000 | 0.05 | 183 |
| Crohn Disease | DOMINO | STRING<br>(401872) | 175000 | 0.05 | 246 |
| Breast Carcinoma | DOMINO | STRING<br>(373074) | 175000 | 0.075 | 527 |
| Schizophrenia | DOMINO | STRING<br>(405088) | 175000 | 0.075 | 315 |

**Table S2:** Results of additional altered subnetwork identification methods on cancer driver mutation *p*-values from TCGA tumor samples from Table 1.

| **Method** | **STRING network** |  |  |  |  |  |  |
| --- | --- | --- | --- | --- | --- | --- | --- |
|  | **Subnetwork size** | **CGC** |  | **OncoKB** |  | **TCGA** |  |
|  |  | *Number* | *F-measure* | *Number* | *F-measure* | *Number* | *F-measure* |
| NetMix2 (connected family) | 266 | 128 | 0.295 | 129 | 0.309 | 144 | 0.534 |
| NetMix | 2256 | 113 | 0.079 | 101 | 0.075 | 38 | 0.03 |
| Heinz (FDR=0.005) | 285 | 130 | 0.293 | 131 | 0.307 | 146 | 0.523 |
| Network-only (degree) | 280 | 8 | 0.018 | 8 | 0.019 | 7 | 0.025 |

## Footnotes

1 A related problem is the identification of altered subnetworks according to network topology alone. Many of the leading methods for this problem were benchmarked in a recent DREAM competition [48].

2 https://gdc.cancer.gov/about-data/publications/pancan-driver

3 https://github.com/shfong/Network-Boosted-GWAS/tree/master/examples/Data

4 https://github.com/Shamir-Lab/EMP/tree/master/data/emp_test/original_datasets

5 https://cancer.sanger.ac.uk/census

6 https://www.oncokb.org/cancerGenes

7 https://www.disgenet.org/downloads

## References

[1] Elena Nabieva, K Jim, A Agarwal, B Chazelle, and M Singh. Whole-proteome prediction of protein function via graph-theoretic analysis of interaction maps. Bioinformatics, 21:i302–i310, 06 2005.

[2] Minghua Deng, Kui Zhang, Shipra Mehta, Ting Chen, and Fengzhu Sun. Prediction of protein function using protein–protein interaction data. Journal of Computational Biology, 10(6):947–960, 2003.

[3] Hon Nian Chua, Wing-kin Sung, and Limsoon Wong. Exploiting indirect neighbours and topological weight to predict protein function from protein-protein interactions. Bioinformatics, 22(13):1623– 1630, 2006.

[4] Roded Sharan, Igor Ulitsky, and Ron Shamir. Network-based prediction of protein function. Molecular Systems Biology, 3:88–88, 2007.

[5] Predrag Radivojac, Wyatt T Clark, Tal Ronnen Oron, Alexandra M Schnoes, Tobias Wittkop, Artem Sokolov, Kiley Graim, et al. A large-scale evaluation of computational protein function prediction. Nature Methods, 10(3):221–227, 2013.

[6] Trey Ideker, O Ozier, B Schwikowski, and Siegel A. F. Discovering regulatory and signalling circuits in molecular interaction networks. Bioinformatics, 18(Suppl 1):S233–S240, 2002.

[7] Igor Ulitsky and Ron Shamir. Identification of functional modules using network topology and high-throughput data. BMC Systems Biology, 1(1):8, 2007.

[8] Marcus T Dittrich, G Klau, A Rosenwald, T Dandekar, and T Muller. Identifying functional modules in protein–protein interaction networks: an integrated exact approach. Bioinformatics, 24(13):i223– i231, 2008.

[9] Alberto de la Fuente. From ‘differential expression’ to ‘differential networking’ – identification of dysfunctional regulatory networks in diseases. Trends in Genetics, 26(7):326–333, 2010.

[10] Dong-Yeon Cho, Yoo-Ah Kim, and Teresa M. Przytycka. Chapter 5: Network biology approach to complex diseases. PLOS Computational Biology, 8(12):1–11, 2012.

[11] Jianguo Xia, Erin E Gill, and Robert E W Hancock. NetworkAnalyst for statistical, visual and network-based meta-analysis of gene expression data. Nature Protocols, 10(6):823–844, 2015.

[12] Sebastian Vlaic, Theresia Conrad, Christian Tokarski-Schnelle, Mika Gustafsson, Uta Dahmen, Reinhard Guthke, and Stefan Schuster. ModuleDiscoverer: Identification of regulatory modules in protein-protein interaction networks. Scientific Reports, 8(1):433, 2018.

[13] Insuk Lee, U Martin Blom, Peggy I Wang, Jung Eun Shim, and Edward M Marcotte. Prioritizing candidate disease genes by network-based boosting of genome-wide association data. Genome Research, 21(7):1109–1121, 2011.

[14] Andrea Califano, Atul J Butte, Stephen Friend, Trey Ideker, and Eric Schadt. Leveraging models of cell regulation and GWAS data in integrative network-based association studies. Nature Genetics, 44(8):841–847, 2012.

[15] Mark DM Leiserson, Jonathan V Eldridge, Sohini Ramachandran, and Benjamin J Raphael. Network analysis of GWAS data. Current Opinion in Genetics & Development, 23(6):602–610, 2013.

[16] Fereydoun Hormozdiari, Osnat Penn, Elhanan Borenstein, and Evan E Eichler. The discovery of integrated gene networks for autism and related disorders. Genome Research, 25(1):142–154, 2015.

[17] Sean Robinson, Jaakko Nevalainen, Guillaume Pinna, Anna Campalans, J Pablo Radicella, and Laurent Guyon. Incorporating interaction networks into the determination of functionally related hit genes in genomic experiments with Markov random fields. Bioinformatics, 33(14):i170–i179, 07 2017.

[18] Justin K Huang, Daniel E Carlin, Michael Ku Yu, Wei Zhang, Jason F Kreisberg, Pablo Tamayo, and Trey Ideker. Systematic evaluation of molecular networks for discovery of disease genes. Cell Systems, 6(4):484–495, 2018.

[19] Rod K Nibbe, Mehmet Koyutürk, and Mark R Chance. An integrative-omics approach to identify functional sub-networks in human colorectal cancer. PLOS Computational Biology, 6(1):e1000639, 2010.

[20] Fabio Vandin, Eli Upfal, and Benjamin J Raphael. Algorithms for detecting significantly mutated pathways in cancer. Journal of Computational Biology, 18(3):507–522, 2011.

[21] Matan Hofree, John P Shen, Hannah Carter, Andrew Gross, and Trey Ideker. Network-based stratification of tumor mutations. Nature Methods, 10(11):1108–1115, 2013.

[22] Pau Creixell, Jüri Reimand, Syed Haider, Guanming Wu, Tatsuhiro Shibata, Miguel Vazquez, Ville Mustonen, Abel Gonzalez-Perez, John Pearson, Chris Sander, Benjamin J Raphael, Debora S Marks, BF Francis Ouellette, Alfonso Valencia, Gary D Bader, Paul C Boutros, Joshua M Stuart, Rune Linding, Nuria Lopez-Bigas, Lincoln D Stein, Mutation Consequences, and Pathway Analysis Working Group of the International Cancer Genome Consortium. Pathway and network analysis of cancer genomes. Nature Methods, 12(7):615–621, 07 2015.

[23] Mark D M Leiserson, Fabio Vandin, Hsin-Ta Wu, Jason R Dobson, et al. Pan-cancer network analysis identifies combinations of rare somatic mutations across pathways and protein complexes. Nature Genetics, 47(2):106–114, 2015.

[24] Raunak Shrestha, Ermin Hodzic, Thomas Sauerwald, Phuong Dao, Kendric Wang, Jake Yeung, Shawn Anderson, Fabio Vandin, Gholamreza Haffari, Colin C Collins, and S Cenk Sahinalp. HIT’nDRIVE: patient-specific multidriver gene prioritization for precision oncology. Genome Research, 27(9):1573–1588, 2017.

[25] modENCODE Consortium, Sushmita Roy, Jason Ernst, Peter V Kharchenko, Pouya Kheradpour, et al. Identification of functional elements and regulatory circuits by drosophila modencode. Science, 330(6012):1787–1797, 2010.

[26] Xin Wang, Camille Terfve, John C Rose, and Florian Markowetz. HTSanalyzeR: an R/Bioconductor package for integrated network analysis of high-throughput screens. Bioinformatics, 27(6):879–880, 01 2011.

[27] Bjarni V Halldórsson and Roded Sharan. Network-based interpretation of genomic variation data. Journal of Molecular Biology, 425(21):3964–3969, 2013.

[28] Bonnie Berger, Jian Peng, and Mona Singh. Computational solutions for omics data. Nature Reviews Genetics, 14(5):333–346, 2013.

[29] Alex J. Cornish and Florian Markowetz. SANTA: Quantifying the functional content of molecular networks. PLOS Computational Biology, 10(9):e1003808.#x2013;, 09 2014.

[30] Vladimir Gligorijević and Nataša Pržulj. Methods for biological data integration: perspectives and challenges. Journal of the Royal Society Interface, 12(112):20150571, 2015.

[31] Jörg Menche, Amitabh Sharma, Maksim Kitsak, Susan Dina Ghiassian, Marc Vidal, Joseph Loscalzo, and Albert-László Barabási. Uncovering disease-disease relationships through the incomplete human interactome. Science, 347(6224):1257601–1257601, 2015.

[32] Susan Dina Ghiassian, Jörg Menche, and Albert-László Barabási. A DIseAse MOdule Detection (DIAMOnD) algorithm derived from a systematic analysis of connectivity patterns of disease proteins in the human interactome. PLOS Computational Biology, 11(4):e1004120.#x2013;, 04 2015.

[33] Deborah Chasman, Alireza Fotuhi Siahpirani, and Sushmita Roy. Network-based approaches for analysis of complex biological systems. Current Opinion in Biotechnology, 39:157 – 166, 2016.

[34] Yunan Luo, Xinbin Zhao, Jingtian Zhou, Jinglin Yang, Yanqing Zhang, Wenhua Kuang, Jian Peng, Ligong Chen, and Jianyang Zeng. A network integration approach for drug-target interaction prediction and computational drug repositioning from heterogeneous information. Nature Communications, 8(1):573, 2017.

[35] Sergio Picart-Armada, Steven J. Barrett, David R. Willé, Alexandre Perera-Lluna, Alex Gutteridge, and Benoit H. Dessailly. Benchmarking network propagation methods for disease gene identification. PLOS Computational Biology, 15(9):1–24, 09 2019.

[36] Koyel Mitra, Anne-Ruxandra Carvunis, Sanath Kumar Ramesh, and Trey Ideker. Integrative approaches for finding modular structure in biological networks. Nature Reviews Genetics, 14(10):719– 732, 2013.

[37] Peilin Jia and Zhongming Zhao. Network assisted analysis to prioritize GWAS results: principles, methods and perspectives. Human Genetics, 133(2):125–138, 02 2014.

[38] Lenore Cowen, Trey Ideker, Benjamin J. Raphael, and Roded Sharan. Network propagation: a universal amplifier of genetic associations. Nature Reviews Genetics, 18(9):551–562, 2017.

[39] Christos M. Dimitrakopoulos and Niko Beerenwinkel. Computational approaches for the identification of cancer genes and pathways. WIREs Systems Biology and Medicine, 9(1):e1364, 2017.

[40] Jason Weston, Andre Elisseeff, Dengyong Zhou, Christina S. Leslie, and William Stafford Noble. Protein ranking: From local to global structure in the protein similarity network. Proceedings of the National Academy of Sciences, 101(17):6559–6563, 2004.

[41] Sebastian Köhler, Sebastian Bauer, Denise Horn, and Peter N. Robinson. Walking the interactome for prioritization of candidate disease genes. The American Journal of Human Genetics, 82(4):949–958, 2008.

[42] Lawrence Page, Sergey Brin, Rajeev Motwani, and Terry Winograd. The PageRank citation ranking: Bringing order to the web. Technical Report 1999-66, Stanford InfoLab, November 1999.

[43] Dengyong Zhou, Olivier Bousquet, Thomas Lal, Jason Weston, and Bernhard Schölkopf. Learning with local and global consistency. In Advances in Neural Information Processing Systems, volume 16. MIT Press, 2004.

[44] Mengfei Cao, Hao Zhang, Jisoo Park, Noah M. Daniels, Mark E. Crovella, Lenore J. Cowen, and Benjamin Hescott. Going the distance for protein function prediction: A new distance metric for protein interaction networks. PLOS ONE, 8(10):1–12, 10 2013.

[45] Lenore Cowen, Kapil Devkota, Xiaozhe Hu, James M. Murphy, and Kaiyi Wu. Diffusion state distances: Multitemporal analysis, fast algorithms, and applications to biological networks. SIAM Journal on Mathematics of Data Science, 3(1):142–170, 2021.

[46] Fabio Vandin, Eli Upfal, and Benjamin J Raphael. De novo discovery of mutated driver pathways in cancer. Genome Research, 22(2):375–385, 2012.

[47] Isabel M. Kloumann, Johan Ugander, and Jon Kleinberg. Block models and personalized PageRank. Proceedings of the National Academy of Sciences, 114(1):33–38, 2017.

[48] Sarvenaz Choobdar, Mehmet E Ahsen, Jake Crawford, Mattia Tomasoni, Tao Fang, David Lamparter, Junyuan Lin, Benjamin Hescott, Xiaozhe Hu, Johnathan Mercer, et al. Assessment of network module identification across complex diseases. Nature Methods, 16(9):843–852, 2019.

[49] Trey Ideker, Vesteinn Thorsson, Jeffrey A Ranish, Rowan Christmas, Jeremy Buhler, Jimmy K Eng, Roger Bumgarner, David R Goodlett, Ruedi Aebersold, and Leroy Hood. Integrated genomic and proteomic analyses of a systematically perturbed metabolic network. Science, 292(5518):929–934, 2001.

[50] Chloé-Agathe Azencott, Dominik Grimm, Mahito Sugiyama, Yoshinobu Kawahara, and Karsten M. Borgwardt. Efficient network-guided multi-locus association mapping with graph cuts. Bioinformatics, 29(13):i171–i179, 06 2013.

[51] Yuanlong Liu, Myriam Brossard, Damian Roqueiro, Patricia Margaritte-Jeannin, Chloé Sarnowski, Emmanuelle Bouzigon, and Florence Demenais. SigMod: an exact and efficient method to identify a strongly interconnected disease-associated module in a gene network. Bioinformatics, 33(10):1536– 1544, 01 2017.

[52] Ery Arias-Castro, David L. Donoho, and Xiaoming Huo. Adaptive multiscale detection of filamentary structures in a background of uniform random points. The Annals of Statistics, 34(1):326–349, 2006.

[53] Ery Arias-Castro, Emmanuel J. Candès, Hannes Helgason, and Ofer Zeitouni. Searching for a trail of evidence in a maze. The Annals of Statistics, 36(4):1726–1757, 2008.

[54] Ery Arias-Castro Emmanuel, J. Candès, and Arnaud Durand. Detection of an anomalous cluster in a network. The Annals of Statistics, 39(1):278–304, 2011.

[55] James Sharpnack, Akshay Krishnamurthy, and Aarti Singh. Near-optimal anomaly detection in graphs using Lovász extended scan statistic. In Proceedings of the 26th International Conference on Neural Information Processing Systems Volume 2, NIPS’13, page 1959–1967, 2013.

[56] James Sharpnack, Aarti Singh, and Alessandro Rinaldo. Changepoint detection over graphs with the spectral scan statistic. In Artificial Intelligence and Statistics, pages 545–553, 2013.

[57] Louigi Addario-Berry, Nicolas Broutin, Luc Devroye, and Gábor Lugosi. On combinatorial testing problems. The Annals of Statistics, 38(5):3063–3092, 2010.

[58] James Sharpnack, Alessandro Rinaldo, and Aarti Singh. Detecting anomalous activity on networks with the graph fourier scan statistic. IEEE Transactions on Signal Processing, 64(2):364–379, 2016.

[59] Iryna Nikolayeva, Oriol Guitart Pla, and Benno Schwikowski. Network module identification–a widespread theoretical bias and best practices. Methods, 132:19–25, 2018.

[60] Matthew A Reyna, Uthsav Chitra, Rebecca Elyanow, and Benjamin J Raphael. NetMix: A network-structured mixture model for reduced-bias estimation of altered subnetworks. Journal of Computational Biology, 28(5):469–484, 2021.

[61] Uthsav Chitra, Kimberly Ding, Jasper C.H. Lee, and Benjamin J Raphael. Quantifying and reducing bias in maximum likelihood estimation of structured anomalies. In Proceedings of the 38th International Conference on Machine Learning, pages 1908–1919. PMLR, 18–24 Jul 2021.

[62] Oron Vanunu, O Magger, E Ruppin, T Shlomi, and R Sharan. Associating genes and protein complexes with disease via network propagation. PLOS Computational Biology, 6(1):e1000641, 2010.

[63] Fabio Vandin, Patrick Clay, Eli Upfal, and Benjamin J Raphael. Discovery of mutated subnetworks associated with clinical data in cancer. In Pacific Symposium on Biocomputing, volume 17, pages 55–66, 2012.

[64] Matthew A Reyna, Mark DM Leiserson, and Benjamin J Raphael. Hierarchical HotNet: identifying hierarchies of altered subnetworks. Bioinformatics, 34(17):i972–i980, 2018.

[65] Evan O Paull, Daniel E Carlin, Mario Niepel, Peter K Sorger, David Haussler, and Joshua M Stuart. Discovering causal pathways linking genomic events to transcriptional states using Tied Diffusion Through Interacting Events (TieDIE). Bioinformatics, 29(21):2757–2764, 2013.

[66] Gal Barel and Ralf Herwig. NetCore: a network propagation approach using node coreness. Nucleic Acids Research, 48(17):e98–e98, 07 2020.

[67] Olga Lazareva, Jan Baumbach, Markus List, and David B Blumenthal. On the limits of active module identification. Briefings in Bioinformatics, 22(5), 03 2021.

[68] Hagai Levi, Ran Elkon, and Ron Shamir. DOMINO: a network-based active module identification algorithm with reduced rate of false calls. Molecular Systems Biology, 17(1):e9593, 2021.

[69] Bradley Efron. Large-scale simultaneous hypothesis testing: the choice of a null hypothesis. Journal of the American Statistical Association, 99(465):96–104, 2004.

[70] Bradley Efron. Correlation and large-scale simultaneous significance testing. Journal of the American Statistical Association, 102(477):93–103, 2007.

[71] Bradley Efron. Size, power and false discovery rates. The Annals of Statistics, 35(4):1351–1377, 2007.

[72] Martin Kulldorff. A spatial scan statistic. Communications in Statistics-Theory and Methods, 26(6):1481–1496, 1997.

[73] Joseph Glaz, Joseph Naus, and Sylvan Wallenstein. Scan Statistics. Springer-Verlag New York, 2001.

[74] Jose Cadena, Feng Chen, and Anil Vullikanti. Near-optimal and practical algorithms for graph scan statistics with connectivity constraints. ACM Transactions on Knowledge Discovery from Data, 13(2):20:1–20:33, 2019.

[75] Wei Pan, Jizhen Lin, and Chap T. Le. A mixture model approach to detecting differentially expressed genes with microarray data. Functional & Integrative Genomics, 3(3):117–124, 2003.

[76] G.J. McLachlan, R. W. Bean, and L. Ben-Tovim Jones. A simple implementation of a normal mixture approach to differential gene expression in multiclass microarrays. Bioinformatics, 22(13):1608–1615, 2006.

[77] David Donoho and Jiashun Jin. Higher criticism for detecting sparse heterogeneous mixtures. The Annals of Statistics, 32(3):962–994, 2004.

[78] T. Tony Cai, Jiashun Jin, and Mark G. Low. Estimation and confidence sets for sparse normal mixtures. The Annals of Statistics, 35(6):2421–2449, 2007.

[79] Stan Pounds and Stephan W Morris. Estimating the occurrence of false positives and false negatives in microarray studies by approximating and partitioning the empirical distribution of p-values. Bioinformatics, 19(10):1236–1242, 2003.

[80] Zheng Guo, Yongjin Li, Xue Gong, Chen Yao, Wencai Ma, Dong Wang, Yanhui Li, Jing Zhu, Min Zhang, D. Yang, and Jing Wang. Edge-based scoring and searching method for identifying condition-responsive protein–protein interaction sub-network. Bioinformatics, 23(16):2121–2128, 2007.

[81] Chloé-Agathe Azencott, Dominik Grimm, Mahito Sugiyama, Yoshinobu Kawahara, and Karsten M. Borgwardt. Efficient network-guided multi-locus association mapping with graph cuts. Bioinformatics, 29(13):i171–i179, 06 2013.

[82] Marzieh Ayati, Sinan Erten, Mark R. Chance, and Mehmet Koyutürk. MOBAS: identification of disease-associated protein subnetworks using modularity-based scoring. EURASIP Journal on Bioinformatics and Systems Biology, 2015(1):7, 2015.

[83] Gurobi Optimization, LLC. Gurobi Optimizer Reference Manual, 2021.

[84] Jishnu Das and Haiyuan Yu. HINT: High-quality protein interactomes and their applications in understanding human disease. BMC Systems Biology, 6(1):92, 2012.

[85] Thomas Rolland, Murat Taşan, Benoit Charloteaux, Samuel J Pevzner, Quan Zhong, Nidhi Sahni, Song Yi, Irma Lemmens, Celia Fontanillo, Roberto Mosca, Atanas Kamburov, Susan D Ghiassian, Xinping Yang, Lila Ghamsari, Dawit Balcha, Bridget E Begg, Pascal Braun, Marc Brehme, Martin P Broly, Anne-Ruxandra Carvunis, Dan Convery-Zupan, Roser Corominas, Jasmin Coulombe-Huntington, Elizabeth Dann, Matija Dreze, Amélie Dricot, Changyu Fan, Eric Franzosa, Fana Gebreab, Bryan J Gutierrez, Madeleine F Hardy, Mike Jin, Shuli Kang, Ruth Kiros, Guan Ning Lin, Katja Luck, Andrew MacWilliams, Jörg Menche, Ryan R Murray, Alexandre Palagi, Matthew M Poulin, Xavier Rambout, John Rasla, Patrick Reichert, Viviana Romero, Elien Ruyssinck, Julie M Sahalie, Annemarie Scholz, Akash A Shah, Amitabh Sharma, Yun Shen, Kerstin Spirohn, Stanley Tam, Alexander O Tejeda, Shelly A Trigg, Jean-Claude Twizere, Kerwin Vega, Jennifer Walsh, Michael E Cusick, Yu Xia, Albert-László Barabási, Lilia M Iakoucheva, Patrick Aloy, Javier De Las Rivas, Jan Tavernier, Michael A Calderwood, David E Hill, Tong Hao, Frederick P Roth, and Marc Vidal. A proteome-scale map of the human interactome network. Cell, 159(5):1212–1226, 2014.

[86] Heiko Horn, Michael S Lawrence, Candace R Chouinard, Yashaswi Shrestha, Jessica Xin Hu, et al. NetSig: network-based discovery from cancer genomes. Nature Methods, 15(1):61–66, 01 2018.

[87] Michael S Lawrence, Petar Stojanov, Craig H Mermel, James T Robinson, Levi A Garraway, Todd R Golub, Matthew Meyerson, Stacey B Gabriel, Eric S Lander, and Gad Getz. Discovery and saturation analysis of cancer genes across 21 tumour types. Nature, 505(7484):495–501, 2014.

[88] Matthew H Bailey, Collin Tokheim, Eduard Porta-Pardo, Sohini Sengupta, Denis Bertrand, Amila Weerasinghe, Antonio Colaprico, Michael C Wendl, Jaegil Kim, Brendan Reardon, et al. Comprehensive characterization of cancer driver genes and mutations. Cell, 173(2):371–385, 2018.

[89] Damian Szklarczyk, Andrea Franceschini, Stefan Wyder, Kristoffer Forslund, Davide Heller, Jaime Huerta-Cepas, Milan Simonovic, Alexander Roth, Alberto Santos, Kalliopi P Tsafou, et al. STRING v10: protein–protein interaction networks, integrated over the tree of life. Nucleic Acids Research, 43(D1):D447–D452, 2015.

[90] John G Tate, Sally Bamford, Harry C Jubb, Zbyslaw Sondka, David M Beare, Nidhi Bindal, Harry Boutselakis, Charlotte G Cole, Celestino Creatore, Elisabeth Dawson, et al. COSMIC: the catalogue of somatic mutations in cancer. Nucleic acids research, 47(D1):D941–D947, 2019.

[91] Debyani Chakravarty, Jianjiong Gao, Sarah Phillips, Ritika Kundra, Hongxin Zhang, Jiaojiao Wang, Julia E Rudolph, Rona Yaeger, Tara Soumerai, Moriah H Nissan, et al. OncoKB: a precision oncology knowledge base. JCO Precision Oncology, 1:1–16, 2017.

[92] AI Velghe, S Van Cauwenberghe, AA Polyansky, Damini Chand, CP Montano-Almendras, S Charni, Bengt Hallberg, Ahmed Essaghir, and Jean-Baptiste Demoulin. PDGFRA alterations in cancer: characterization of a gain-of-function V536E transmembrane mutant as well as loss-of-function and passenger mutations. Oncogene, 33(20):2568–2576, 2014.

[93] Sebastiano Battaglia, Orla Maguire, and Moray J Campbell. Transcription factor co-repressors in cancer biology: roles and targeting. International Journal of Cancer, 126(11):2511–2519, 2010.

[94] Daniel E. Carlin, Samson H. Fong, Yue Qin, Tongqiu Jia, Justin K. Huang, Bokan Bao, Chao Zhang, and Trey Ideker. A fast and flexible framework for network-assisted genomic association. iScience, 16:155–161, 2019.

[95] Janet Piñero, Juan Manuel Ramírez-Anguita Josep Saüch-Pitarch, Francesco Ronzano, Emilio Centeno, Ferran Sanz, and Laura I Furlong. The DisGeNET knowledge platform for disease genomics: 2019 update. Nucleic Acids Research, 48(D1):D845–D855, 2020.

[96] Casey S Greene, Arjun Krishnan, Aaron K Wong, Emanuela Ricciotti, Rene A Zelaya, Daniel S Himmelstein, Ran Zhang, Boris M Hartmann, Elena Zaslavsky, Stuart C Sealfon, Daniel I Chasman, Garret A FitzGerald, Kara Dolinski, Tilo Grosser, and Olga G Troyanskaya. Understanding multicellular function and disease with human tissue-specific networks. Nature Genetics, 47(6):569–576, 2015.

[97] Samson H Fong, Daniel E Carlin, Kivilcim Ozturk, 2018 UCSD Network Biology Class, and Trey Ideker. Strategies for network GWAS evaluated using classroom crowd science. Cell systems, 8(4):275–280, 04 2019.

[98] Jesse Davis and Mark Goadrich. The relationship between precision-recall and roc curves. In Proceedings of the 23rd International Conference on Machine Learning, ICML ‘06, page 233–240, New York, NY, USA, 2006. Association for Computing Machinery.

[99] David Lamparter, Daniel Marbach, Rico Rueedi, Zoltán Kutalik, and Sven Bergmann. Fast and rigorous computation of gene and pathway scores from SNP-based summary statistics. PLOS Computational Biology, 12(1):e1004714, 2016.

[100] Priyanka Nakka, Benjamin J Raphael, and Sohini Ramachandran. Gene and network analysis of common variants reveals novel associations in multiple complex diseases. Genetics, 204(2):783–798, 2016.

[101] Emil Uffelmann, Qin Qin Huang, Nchangwi Syntia Munung, Jantina de Vries, Yukinori Okada, Alicia R Martin, Hilary C Martin, Tuuli Lappalainen, and Danielle Posthuma. Genome-wide association studies. Nature Reviews Methods Primers, 1(1):1–21, 2021.

[102] Xingyi Guo, Weiqiang Lin, Jiandong Bao, Qiuyin Cai, Xiao Pan, Mengqiu Bai, Yuan Yuan, Jiajun Shi, Yaqiong Sun, Mi-Ryung Han, et al. A comprehensive cis-eQTL analysis revealed target genes in breast cancer susceptibility loci identified in genome-wide association studies. The American Journal of Human Genetics, 102(5):890–903, 2018.

[103] Zhihong Zhu, Futao Zhang, Han Hu, Andrew Bakshi, Matthew R Robinson, Joseph E Powell, Grant W Montgomery, Michael E Goddard, Naomi R Wray, Peter M Visscher, et al. Integration of summary data from GWAS and eQTL studies predicts complex trait gene targets. Nature Genetics, 48(5):481–487, 2016.

[104] Evan A Boyle, Yang I Li, and Jonathan K Pritchard. An expanded view of complex traits: from polygenic to omnigenic. Cell, 169(7):1177–1186, 2017.

[105] Bradley Efron, Brit B Turnbull, and Balasubramanian Narasimhan. locfdr: Computes local false discovery rates. R package version, 1:1–7, 2011.

[106] Michael S. Lawrence, Petar Stojanov, Craig H. Mermel, James T. Robinson, Levi A. Garraway, Todd R. Golub, Matthew Meyerson, Stacey B. Gabriel, Eric S. Lander, and Gad Getz. Discovery and saturation analysis of cancer genes across 21 tumour types. Nature, 505(7484):495, 2014.

[107] Yoav Benjamini and Yosef Hochberg. Controlling the false discovery rate: a practical and powerful approach to multiple testing. Journal of the Royal Statistical Society: Series B (Methodological), 57(1):289–300, 1995.

[108] Michele Carbone, Haining Yang, Harvey I Pass, Thomas Krausz, Joseph R Testa, and Giovanni Gaudino. BAP1 and cancer. Nature Reviews Cancer, 13(3):153–159, 2013.

